# There is an “I” in team: Intrapersonal physiological synchrony complements interpersonal physiological synchrony in team interaction

**DOI:** 10.64898/2026.09.28.755043

**Authors:** Chen Erez, Sebastian Wallot, Ilanit Gordon

**Author notes:** Corresponding author: Ilanit Gordon, Department of Psychology and Gonda Multidisciplinary Brain Research Center, Bar-Ilan University, Ramat Gan, Israel.

## Abstract

Interpersonal physiological synchrony (IPS) between interaction partners has been studied as a marker of social connectedness and coordination. However, little is known about how such interpersonal synchrony relates to the intrapersonal coordination of individuals’ own autonomic systems during team interaction. The present study examined whether IPS and intrapersonal physiological synchrony (IAPS), defined as the within-person coordination of heart rate (HR), respiration, and electrodermal activity (EDA), function as competing or complementary regulatory processes. It also examined whether IAPS is state-or trait-like and whether teams high or low in both IPS and IAPS had better outcomes. Fifty-five newly formed triads (*N* = 165; *M_age_* = 23.87, 77% female) participated in a laboratory interaction including passive shared experiences and active collaborative tasks while continuous physiological activity was recorded. Synchrony within individuals and between team members was quantified using multidimensional recurrence quantification analysis. Across interaction segments, IAPS was positively associated with IPS in HR, EDA, and respiration. The HR association did not differ across segments, whereas segment differences in the EDA and respiration associations were specification-sensitive. These findings contradict a trade-off hypothesis and instead support a synergistic relationship between IAPS and IPS. IAPS showed both state-and trait-like features: higher levels in passive than active contexts and relative stability across similar contexts. IAPS was not reliably associated with individual-difference measures. Exploratory profile analyses found differences in perceived group efficacy across IAPS/IPS-BPM profiles, although the pattern was not specific to the high-high subgroup and was based on small, unequal subgroups. The IPS-EDA profile test for performance did not survive correction. Together, these results suggest that intrapersonal and interpersonal physiological synchrony coexist as complementary patterns of physiological organization during team interaction. They further position IAPS as a dynamic component of social functioning that merits investigation alongside interpersonal processes.

## Introduction

Physiological autonomic activity is affected by internal physiological demands and our social environment. It is also often temporally coordinated between individuals during social interactions (e.g., coordinated temporal changes in heartbeat or EDA; Butler, 2015; Helm et al., 2012; Palumbo et al., 2017). In the past, interpersonal physiological synchrony (IPS) has been associated with bonding, empathy, coregulation, and joint performance (Levenson & Gottman, 1983; Marci et al., 2007; Palumbo et al., 2017), corresponding with models of interpersonal emotional regulation that emphasize dynamic reciprocal adjustments in partners’ affective and physiological states (Butler & Randall, 2013).

Synchrony in different autonomic modalities might index distinct affective and regulatory processes and thus indicate different interpersonal dynamics (Gordon et al., 2025). Synchrony in heart rate (or interbeat intervals), which is dynamically regulated by parasympathetic and sympathetic activation, is often associated with affiliation and engagement (Feldman et al., 2011). EDA is a marker of sympathetic activity uniquely, more closely associated with arousal, vigilance, or stress (Boucsein et al., 2012; Dawson et al., 2017) and yet EDA synchrony can also be positively associated with relational outcomes (Mayo et al., 2021). Respiration is somewhat different because breathing is primarily regulated automatically but also changes with emotional state and can be intentionally modified. Respiratory synchrony may be related to affective and regulatory demands, yet it may also arise through coordinated speech, turn-taking, or shared task rhythms (Homma & Masaoka, 2008; McFarland, 2001; Müller & Lindenberger, 2011). Taken together, these findings suggest that modality-specific synchrony reflects a mixture of motivational and regulatory processes shaped by task demands, individual differences and social context (Gordon et al., 2025; Mayo & Gordon, 2020; Tomashin et al., 2022).

Despite extensive work on IPS, especially in the last two decades, little is known about how interpersonal synchronization relates to, or is affected by, the internal organization of individuals’ autonomic systems. Central models of autonomic activity, such as neurovisceral integration and polyvagal theory, posit that autonomic subsystems (within the individual) covary in ways that support regulatory efficiency, flexibility in addressing changing demands from the environment, and adaptive behavior (Porges, 2011; Thayer & Lane, 2000). Such physiological coordination may form part of self-regulation, understood more broadly as the set of processes that adaptively adjust mental and physiological states or behavior to context, but it is not equivalent to self-regulation itself (Nigg, 2017). Studies have found, for example, that coupling between phasic EDA and HR accelerations during psychological testing is negatively associated with depression, and positively with task engagement (Kettunen et al., 1998). Other studies have demonstrated that the strength of covariation between parasympathetic heart-rate variability and EDA varies with context and regulatory demands, being higher during rest and weaker or even decoupled during certain tasks (Callara et al., 2021). More broadly, network physiology conceptualizes the organism as a network of interacting physiological systems whose patterns of coupling reorganize across physiological states (Bashan et al., 2012). Some modalities are also linked through identifiable physiological mechanisms, including cardiorespiratory interactions between respiration and cardiac activity (Elstad et al., 2018). Goheen et al. (2024) further describe how brain and breathing activity can synchronize over shorter timescales and attune to the temporal structure of environmental input over longer timescales. However, the issue of how multiple autonomic subsystems covary within individuals during social interaction is underexplored. We refer to this within-person covariation as intrapersonal physiological synchrony (IAPS), defined as the temporal coordination among heart rate, respiration, and EDA. IAPS is therefore a descriptive, multivariate measure of within-person physiological coupling, rather than a direct measure of self-regulatory capacity or evidence of a single underlying physiological mechanism.

One central issue is whether interpersonal and intrapersonal synchrony support or interfere with one another. That is, are they competing, complementary, overlapping, or even unrelated processes? One possibility is a trade-off, in which they each draw resources needed for the other. Support for this view comes from findings that increased interpersonal synchrony can enhance affiliation while impairing self-regulation (Galbusera et al., 2019), and that interpersonal respiratory entrainment during interaction alters individuals’ cardiorespiratory synchrony (Konvalinka et al., 2026). It is also supported by models proposing that attentional demands reduce the stability of internal regulatory processes (Dikker et al., 2017) and the precision of bodily representations (Critchley & Garfinkel, 2018). An alternative ‘synergy’ perspective predicts that IAPS and IPS would co-occur. Vagal tone, or Respiratory Sinus Arrhythmia, is itself a measure indicative of integration between several physiological (cardiorespiratory) functions within the individual (Porges, 2007) and thereof its association with interpersonal coordination (e.g., Lunkenheimer et al., 2016) is indicative of potential synergy. Within this view, individuals with higher IAPS may show stronger IPS, reflecting greater regulatory flexibility.

Fluctuations in IAPS, and its dynamics with IPS may depend on contextual regulatory demands. Contemporary work demonstrated that (Gordon et al., 2025; Mayo & Gordon, 2020, 2025) social situations contain different balances (as well as dynamic shifts) between two computing forces – to coordinate vs. to segregate. The balance between these motivations, which is affected by both social setting and individual differences (e.g., emotion contagion, task-orientation) affects the extent to which IPS is beneficial for a specific interaction, which can also vary as a function of physiological modality.

Similarly, we suggest that different task demands, specifically the level of passive or active engagement with the social situation, will impact the balance between IAPS and IPS. Recent work found that the effect of team demographic composition on IPS could change between passive, mere-copresence and active task collaboration (Ohayon et al., 2026). However, passive shared experiences may evoke synchrony with minimal social and regulatory interference (e.g., exposure to similar stimuli), whereas interactive joint action requires continuous prediction, joint attention, and monitoring of actions (Dikker et al., 2017; Gordon et al., 2025; Mayo & Gordon, 2020), potentially altering the balance between inter-and intrapersonal synchrony. Hence we focus on comparing passive and active interaction stages.

Finally, the balance between IAPS and IPS could have task-relevant implications. As postulated (de Jong et al., 2014; Felps et al., 2006), teams’ interpersonal affective dynamics can result in team members becoming more focused on maintaining self-regulation and distance from the team, affecting team functioning and outcomes. Two central outcomes for team functioning are task performance and perceived group efficacy. Task performance embodies the objective success of the team while team efficacy (also “collective efficacy”) refers to the team’s subjective shared belief in its ability to reach its goals and produce effects by collective action (Bandura, 2000). We thus aim to explore how the balance between IAPS and IPS is associated with team performance and efficacy.

### The Current Study

In this study, we tested how IPS and IAPS unfold dynamically across passive and active phases of social interaction. We first focused on testing competing hypotheses: whether there is a trade-off or synergy between IAPS and IPS. In addition, we focused on IAPS as a construct, and the degree to which it is a stable individual difference or more context-dependent. Lastly, we explored whether teams of specific IAPS-IPS profiles function better.

To that end, we assembled participants into newly formed triads for a laboratory-based, videotaped, and physiologically monitored interaction. We calculated their individual and shared autonomic synchrony throughout the different stages of this interaction. Using synchrony measures of heart rate, EDA, and respiration, we quantified IPS between partners and IAPS within individuals. IAPS was defined as within-person synchrony among heart rate, respiration, and EDA. IPS was quantified separately as synchrony among team members’ heart rates, EDA, and respiration signals. We tested the following hypotheses:

#### Hypothesis 1: competing trade-off and synergy predictions

We tested whether IPS was negatively associated with IAPS across interaction segments, consistent with a trade-off account, or positively related to IAPS, consistent with a synergy account. This was evaluated separately for heart-rate IPS (IPS-BPM), electrodermal IPS (IPS-EDA), and respiration (IPS-RESP).

#### Hypothesis 2: contextual variation in IAPS-IPS coupling

We also tested whether IAPS-IPS associations vary across social interaction contexts. Specifically, we tested whether they differ across the four study segments: baseline, Task 1, affect manipulation, and Task 2.

Robustness analyses for Hypotheses 1 and 2 separately adjusted the IPS-BPM model for segment-specific mean heart rate and the IPS-EDA model for tonic skin conductance level, and all three IPS models for respiratory sinus arrhythmia, triad mean age, pre-task positive and negative affect, and team emotion-regulation composition.

#### Hypothesis 3: context dependence and stability of IAPS (“state or trait”)

We explored the context-dependency of IAPS (i.e., state vs trait) by testing (a) whether participant-level IAPS differed across segments, (b) whether IAPS levels showed cross-segment relative stability (e.g., participants with high IAPS in one segment having high IAPS across all segments), and (c) whether IAPS was associated with more stable self-reported personality and individual-difference measures or differed across established symptom-threshold groups.

#### Hypothesis 4 (exploratory): synchrony profiles and team outcomes

Lastly, we tested whether task-related combinations of high and low IAPS with high and low IPS-BPM or IPS-EDA were associated with objective team performance and perceived group efficacy.

### Method

The project aimed at addressing the interplay between IAPS and IPS was not preregistered. However, this project utilizes data from a larger parent study exploring team composition and dynamics, which was pre-registered here: https://doi.org/10.17605/OSF.IO/PWRZD.

### Participants

Adults with no self-reported history of dyslexia, intellectual disability, or cardiovascular conditions were recruited. The dyslexia and intellectual-disability criteria were relevant to the word-assembly tasks, and the cardiovascular criterion was relevant to physiological measurement. Participants filled out a short survey for entering a lottery for $150 (and course credit, when applicable) and to determine eligibility for a laboratory study offering an additional $30 payment (or additional course credit). A total of 1520 individuals filled out this screening survey. We recruited individuals with average (28.3% of the sample) and low (18.2%) emotion-regulation (ER) capabilities (Erez & Gordon, 2025) and intentionally formed triads with different ER compositions due to the aims of the larger parent study. Group ER composition was not central to the current research questions and hence was examined only as a secondary predictor of IAPS and as an additive covariate in robustness models of the IAPS–IPS associations.

The initial sample included 84 triads (*N* = 252). As this study involved the relatively novel construct of IAPS, we opted to avoid data imputations and retained only teams with complete physiological data. Twenty-nine triads were excluded: 20 with at least one participant with corrupted respiratory data, 8 with corrupted EDA data, and 1 with completely missing physiological data due to computer failure. This resulted in a final sample of 55 triads (*N* = 165) with complete physiological datasets. The sample was predominantly female (77%), single (98.2%), undergraduate students (86.1%), with a mean age of 23.87 (*SD* = 4.67). Socioeconomic status was normally distributed. Excluded teams were significantly older (26.41; *p* <.05), largely due to four excluded triads that each contained one much older participant (50s–60s). Triads were formed according to the parent study’s recorded ER-composition assignment, without deliberate gender matching or balancing. Of the 55 triads, 2 included no women, 8 included one woman, 16 included two women, and 29 included three women (Table S1).

### Materials and Procedure

#### Pre-Study Self-Report Questionnaires

See Figure 1 for visualization of the procedure and central materials. Data collection took place between November 2021 and June 2023. Individuals were invited via social media and university platforms to fill a 10-minute online screening battery on individual differences. In efforts to gather a diverse sample, we routinely posted ads in general social media forums unrelated to academia. The screening battery included the Big Five Inventory (John et al., 1991), Difficulties in Emotion Regulation Scale (DERS; (Gratz & Roemer, 2004), Social Phobia Inventory (Connor et al., 2000), Generalized Anxiety Disorder-7 (GAD-7; Spitzer et al., 2006), the Patient Health Questionnaire-2 assessing depression (PHQ-2; Arroll et al., 2010), and the Single-item Life Satisfaction Scale (Jovanović & Lazić, 2020). Those who scored between 65 and 85, or 95 or above on the DERS met the criteria for average and low ER capabilities, respectively (Erez & Gordon, 2025) and were invited via email to partake in a laboratory study on behavioral and physiological processes during group interaction.

**Figure 1.**
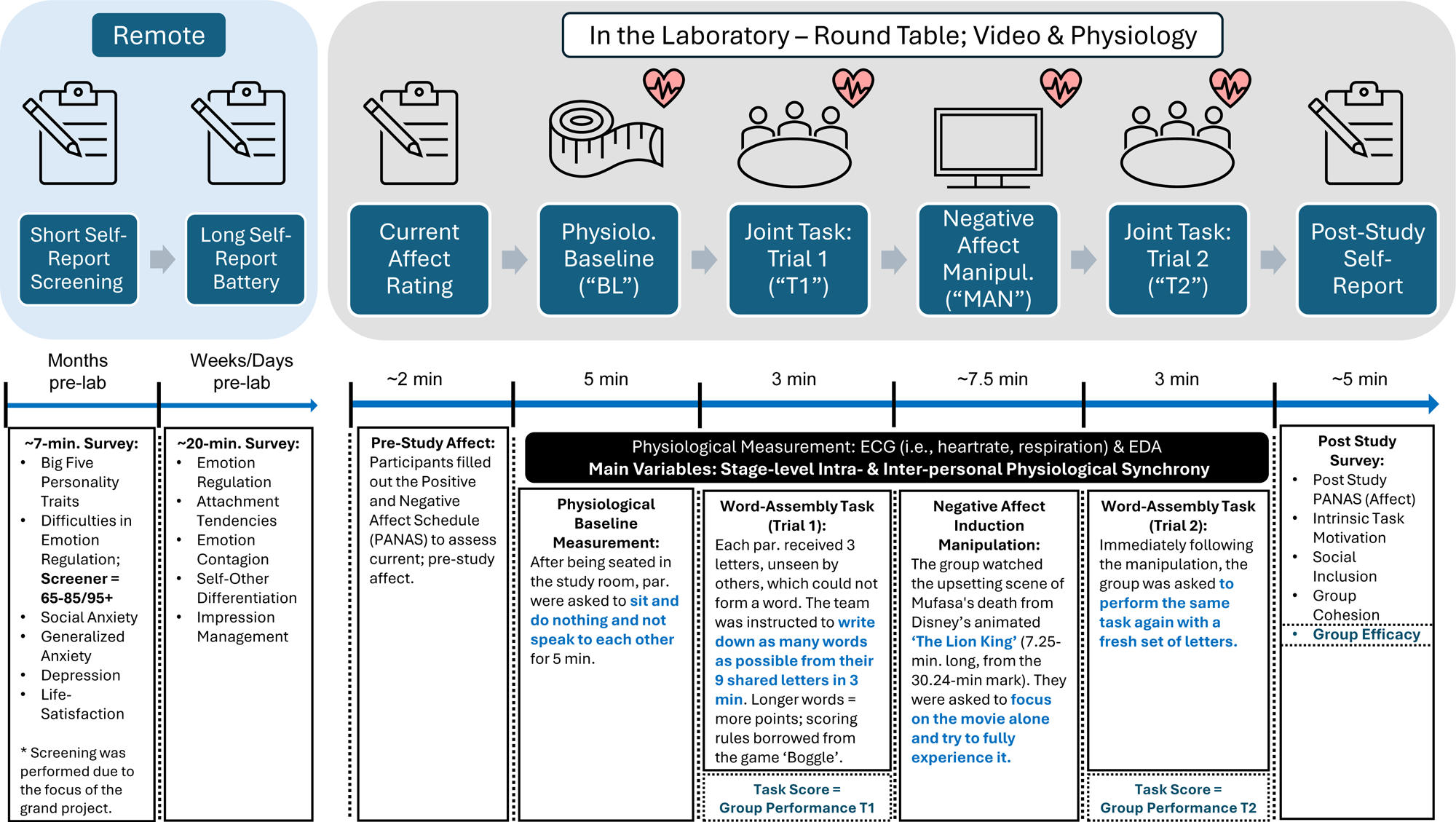
Graphical depiction of study procedure. Participants first filled out a short screening survey. Those who met the criteria were invited to the laboratory group study. Before arriving, they filled out another survey (remotely). In the lab they filled out the PANAS, underwent a baseline measurement, group word-assembly task, watched an upsetting movie scene from The Lion King, had a second word-assembly task followed by a post-study survey.

Those who decided to participate completed an additional 20-minute online survey several days or weeks prior to coming to the lab. They answered the Emotion Contagion scale (Doherty, 1997), Emotion Regulation Questionnaire (Gross & John, 2003), Negative Mood Regulation Scale (Catanzaro & Mearns, 1990), Experiences in Close Relationships-Revised (Mikulincer & Florian, 2000), Differentiation of Self Inventory-Revised (Skowron & Schmitt, 2003), and the Balanced Inventory of Desirable Responding (Paulhus & Reid, 1991).

#### Laboratory Interaction

Participants were instructed to hydrate normally and avoid caffeine and nicotine for a minimum of 2 hours before arriving at the lab. Upon arrival, they washed their palms with water and soap. They were then connected to MindWare Mobile Impedance Cardiograph recording devices (MindWare Technology, Gahanna, OH) in a private room by same-gender experimenters blinded to ER scores and composition. Their physiological activity was continuously monitored.

They were then seated in a videotaped study room (ceiling, and 2 walls) furnished with a round table and a 47“ monitor. Each participant sat across an Ikea VIVALLA tablet stand with a Logitech Brio webcam facing them. They then completed via tablet the Positive and Negative Affect Schedule (PANAS; Watson et al., 1988), modified to include the items “happy” and “sad” due to their relevance to the larger parent study.

The experimental session consisted of four segments: (1) *Physiological Baseline Measurement* (BL: 5 min) - Participants sat quietly and were asked to do nothing and not speak to each other. (2) *Task Trial 1* (T1; 3 min) - A group word-assembly game (rules adapted from “Boggle”). Each member received three unique letters, and the triad was instructed to form as many words as possible using their combined nine letters. By design, each member’s three letters could not form a word, for high-interdependency. Members were instructed to place their letters on their individual tablet stands. They could communicate freely but not physically show their teammates their letters. (3) *Negative Affect Manipulation* (MAN: 7.5 min) - Participants watched an emotionally distressing scene from The Lion King. They were asked to focus on the movie alone and try to fully experience it. The movie was meant to challenge their ER system by generating higher negative affect (mostly sadness) and reducing positive affect (significantly effective in pilot study); following insights from the meta-analysis on emotion induction manipulations by Joseph et al. (2020). A negative affect induction, rather than a neutral or positive induction, was implemented for the purposes of the parent study on ER composition. (4) *Task Trial 2* (T2; 3 min) – A second iteration of the word-assembly game with new letters.

Following T2, participants completed a second PANAS and post-task questionnaires including group cohesion (Podsakoff & MacKenzie, 1994), intrinsic motivation (Ryan, 1982), social inclusion (Aron et al., 1992), and *group efficacy* (Jex & Bliese, 1999) – in which participants indicate their belief in their group’s abilities on 4 items (“I am confident in my team’s ability to perform”) via a 7-point Likert scale (1 = not at all, 7 = very much) (Cronbach’s α = 0.87). To compute group scores, individual self-reports were mean-aggregated at the group level, including group efficacy, one of the central dependent variables in this project. Group overall performance, another central construct in this effort, was calculated by summing each group’s scores on both T1 and T2. One of the 55 physiology-eligible triads lacked complete overall-performance data. It was retained in physiology and efficacy analyses, while analyses involving overall performance used the 54 available complete cases.

#### Physiological Data Acquisition and Preprocessing

Physiological data were acquired at 500 Hz using MindWare Mobile Impedance Cardiographs (MindWare Technology, Gahanna, OH). Electrocardiogram (ECG) was recorded via a modified Lead II configuration. Respiration was derived from impedance cardiography (Z_0_) measured via 4 electrodes placed on the torso. Electrodermal activity (EDA) was recorded via 2 electrodes placed on the thenar and hypothenar eminences of the non-dominant hand which was mostly stationary.

Data were filtered and visually inspected using MindWare analysis software, and artifact correction followed standard guidelines (Boucsein et al., 2012; Quigley et al., 2024). Respiratory data were marked as corrupted if the impedance signal remained flat (i.e., no signal) for more than 10% of the recording duration (analyzed per interaction segment).

#### Calculating Synchrony

IPS and IAPS levels were calculated via Multidimensional Recurrence Quantification Analysis (MdRQA). MdRQA was selected because it allows characterization of nonlinear, multichannel coordination across physiological systems. MdRQA and other recurrence-based analyses have been increasing in usage over the past decade across disciplines for calculating synchrony between multiple time-series (Tomashin et al., 2022; Wenzel et al., 2023; Wiltshire et al., 2022). Unlike univariate or pairwise synchrony measures, MdRQA reconstructs a single phase space from multiple signals, providing an index of shared dynamics (Marwan et al., 2007; Wallot & Leonardi, 2018). This approach aligns with the study’s goal of assessing how interpersonal and intrapersonal autonomic coordination unfold across interaction contexts and whether these forms of coordination support or compete with one another. For IAPS, we assessed coordination among each participant’s heart rate, respiration, and EDA signals. Each group’s IAPS level was calculated by averaging each member’s IAPS level. For IPS, we examined coordination among the three group members’ heart-rate signals, EDA signals, and respiration signals separately. Importantly, IAPS and IPS were treated as distinct but potentially interacting indices rather than statistically orthogonal constructs.

Inter-beat intervals were converted to beats per minute (BPM) and, like respiration (Z_0_), were resampled to 2 Hz to match the lowest sampling rate set by the EDA signal and to ensure temporal alignment across time-series. All data were z-score normalized prior to embedding so that each contributed equally to Euclidean distances in the multidimensional state space (Marwan et al., 2007).

MdRQA parameters were estimated following established procedures (Wallot et al., 2016). Delay (d) and embedding dimension (m) were determined separately for each synchrony analysis (IAPS, IPS-BPM, IPS-EDA, IPS-RESP) across the four study segments (Baseline, T1, Manipulation, T2) before selecting suitable cross-segment parameters for each analysis. That is, for IPS analyses, the focal units for estimation were teams while for IAPS the focal units were individuals. For each analysis, we aimed to use the parameters most suitable for representing the system’s behavior throughout all segments. This approach prioritized comparability across participants, triads, and segments, because differences in %REC could not be produced by unit-specific parameter choices. The trade-off is that the common values may not be optimal for every participant or triad.

Delay was estimated with the multidimensional crqa::mdDelay function, using the first local minimum of the Average Mutual Information (AMI) function for each analytic unit within each segment. The same procedure was used for IAPS and all three IPS modalities. It is generally recommended to use the average delay. For IAPS, the four segment-level mean delays were 8.64, 7.22, 11.53, and 7.21 (mean across segment means = 8.65), and *d* = 8 was selected as a common cross-segment value, to better represent most segments than be elevated too high because of the larger average delay during the Manipulation segment. For IPS-BPM, the corresponding means were 7.36, 7.80, 8.58, and 7.84 (mean = 7.90), yielding *d* = 8. For IPS-EDA, they were 15.62, 10.69, 23.20, and 10.58 (mean = 15.02), yielding *d* = 15. For IPS-RESP, the corresponding means were 2.93, 2.98, 2.87, and 3.00 (mean = 2.95), yielding *d* = 3.

Embedding dimension (*m*) was estimated using the multidimensional crqa::mdFnn function (i.e., the false-nearest-neighbors [FNN] method), and the FNN curves were inspected. For IAPS, the FNN plots showed a low-order solution at *m* = 2 driven by respiratory periodicity and another at *m* = 10. The latter higher-dimensional solution was retained to capture interactions among the three autonomic subsystems. The IPS analyses showed clearer solutions, yielding *m* = 8 for BPM, *m* = 3 for EDA, and *m* = 2 for respiration. All parameter selections were applied consistently across segments for each analysis.

To ensure comparability and adequate representation of system dynamics, a common-radius calibration procedure was used (Webber & Zbilut, 2005). For each analysis, the radius was selected to yield a cross-segment average Recurrence Rate (%REC; used here as the operational synchrony index) of 5%, calibrated to three decimal places. Radius selection was manual and iterative. For each analysis, candidate radii were evaluated until the mean of the four segment-level mean %REC values was approximately 5%. Thus, %REC was not fixed at 5% for each observation; it remained free to vary and served as the sole recurrence-derived outcome (*r* = 4.457 for IAPS, *r* = 4.441 for IPS-BPM, *r* = 1.114 for IPS-EDA, and *r* = 1.325 for IPS-RESP). Additional details regarding the parameter estimation procedure are provided in the “MdRQA Parameter Selection” section of the Supplementary Materials. The sole MdRQA-derived outcome entered into the subsequent analyses was %REC; other recurrence-plot indices were not analyzed.

#### Statistical Analysis

##### Primary Models

The analyses were conducted at the level at which each construct was measured. Participant-level IAPS analyses contained four segment observations nested within participants, with participants nested within triads. IAPS-IPS analyses were conducted at the triad level because IPS is a group-level measure. The final sample included 55 triads and 165 participants, yielding 660 participant-by-segment IAPS observations and 220 triad-by-segment IPS observations.

To test Hypotheses 1 and 2, separate linear mixed models were fitted predicting IPS-BPM, IPS-EDA, and IPS-RESP. Each model included centered triad-level IAPS, segment, and their interaction as fixed effects. The selected IPS-BPM model included uncorrelated triad random intercepts and within-triad IAPS slopes, the selected IPS-EDA model included a triad random intercept, and the selected IPS-RESP model included correlated triad random intercepts and within-triad IAPS slopes. The differences among these random-effects structures arose from applying the same documented maximal-to-simple selection procedure separately to each outcome; the first identifiable, converged, nonsingular candidate meeting any applicable optimizer-stability criterion was retained. Sum-to-zero contrasts were used for segment. Consequently, the IAPS coefficient represented its average association with IPS across the four segments, thus providing findings for the main trade-off versus synergy examination. The omnibus IAPS × segment term tested whether this association differed across segments. Analyses of the segment-specific IAPS slopes and Holm-adjusted comparisons among the six slope pairs further tested the context dependency of the IAPS-IPS association. Twelve same-segment triad-level IAPS-IPS correlations were also reported descriptively and Holm-adjusted as one family, but conclusions were based primarily on the mixed models.

To test Hypothesis 3, participant-level IAPS was predicted as a function of interaction segment, with random intercepts for triad and participant, testing whether IAPS levels fluctuated across the interaction. The omnibus segment test was followed by Holm-adjusted comparisons among the six segment pairs to determine between-segment differences in IAPS levels. In addition, six cross-segment correlations were tested, forming one Holm-adjusted stability family, exploring whether participants’ IAPS remained stable in terms of relative position in the sample. Lastly, to test associations between segment-specific IAPS and self-reported measures that vary in stability (e.g., depression versus extraversion), we calculated 88 Pearson correlations between segment-specific IAPS and the broader individual-difference battery (treated as exploratory and controlled via Benjamini-Hochberg false-discovery-rate procedure). Sixteen two-sided Welch independent-samples *t* tests, Holm-corrected as one family, compared IAPS between participants below versus at or above each established cutoff for ER difficulties, social anxiety, generalized anxiety, and depression.

For the mixed models, raw-scale models are reported in the main text. Interpretations also took into account the additional models described below and reported fully in the Supplementary Materials.

For the exploratory outcome analyses of Hypothesis 4, IAPS, IPS-BPM, and IPS-EDA were averaged across Tasks 1 and 2 and then divided at their sample medians and combined into two composite variables, each defining four IAPS–IPS profiles: high-high, high-low, low-high, and low-low. Objective performance and perceived group efficacy were compared across the four profiles using Welch one-way analyses of variance. Analyses involving objective performance used 54 complete cases; missing performance did not exclude the triad from physiology or efficacy analyses. Holm correction was applied across the four omnibus tests. Games–Howell comparisons, which accommodate unequal variances and cell sizes and intrinsically adjust the six pairwise comparisons, were interpreted only when both the corresponding omnibus test remained significant after Holm correction and the Games–Howell-adjusted pairwise *p* value was below.05.

##### Mixed-Model Fit and Effect Sizes

Following recommendations for transparent mixed-model reporting (Meteyard & Davies, 2020; Schubert et al., 2025), we report marginal and conditional *R²* with 1,000-iteration bootstrap confidence intervals for each final model.

Adjusted ICCs summarized the residual clustering remaining after accounting for the model’s fixed effects; for random-slope models, we averaged the adjusted ICC over the observed within-triad IAPS values. To summarize term-specific effects, we derived approximate partial *R²* and Cohen’s *f²* from Type III Satterthwaite *F* tests. Finally, unstandardized coefficients and 95% confidence intervals served as our primary effect estimates.

##### Model Specification, Diagnostics, and Robustness

We fitted mixed models using restricted maximum likelihood and evaluated a documented maximal-to-simple hierarchy of random-effects structures. For the triad-level IPS models, the hierarchy tested correlated and uncorrelated random segment slopes, followed by correlated and uncorrelated random slopes for within-triad IAPS deviations, and finally a triad random-intercept model. For the participant-level IAPS model, correlated and uncorrelated segment slopes were tested at the triad and participant levels before simplification to random intercepts. We retained the first identifiable, converged, nonsingular candidate; AIC and BIC were descriptive and were not used to select among nonnested candidates. The selected IPS-BPM model included a triad random intercept and an uncorrelated random slope for within-triad IAPS, the IPS-EDA model included a triad random intercept, the IPS-RESP model included a triad random intercept and correlated random slope for within-triad IAPS, and the participant-level IAPS model included random intercepts for triad and participant. Nonsingular correlated IAPS-slope candidates were additionally required to be stable across three optimizers; near-boundary intercept-slope correlations were flagged for review rather than treated automatically as singular.

To evaluate model adequacy, we checked for convergence and singularity, and examined residual-versus-fitted and scale-location plots alongside quantile-quantile (Q-Q) plots for both residuals and random effects. We also conducted supplementary diagnostic checks of residual normality and heteroscedasticity, assessed collinearity, and ran triad-level influence analyses. We did not exclude any observation or triad solely for being flagged by a diagnostic.

We retained raw-scale models for our primary analyses. To evaluate robustness, we tested models that allowed for segment-specific residual variances and, separately, applied log1p transformations to the outcomes. We also used a within-between decomposition to separate each triad’s mean IAPS across segments from its segment-specific deviations around that mean. The between-triad term estimated whether triads with overall higher IAPS had higher IPS while the within-triad term estimated whether segment-to-segment deviations from a triad’s own mean IAPS predicted corresponding changes in IPS. Finally, we refitted all primary models omitting one triad at a time to ensure results were not driven by single influential groups.

To test robustness of the IAPS-IPS associations above and beyond other potentially central factors, we fitted separate models adjusting the IPS-BPM model for segment-specific mean heart rate and the IPS-EDA model for tonic skin conductance level. We also separately tested all three IPS models while including respiratory sinus arrhythmia, triad mean age, and the combination of pre-task positive and negative affect. Team emotion-regulation composition was examined only in secondary analyses and was not included in the primary models. The secondary participant-level IAPS model included ER composition, segment, and their interaction, allowing overall differences among recorded compositions and possible differences in their segment patterns to be tested separately. ER composition was subsequently included as an additive covariate in the IAPS–IPS robustness models. Gender composition was summarized descriptively and was not included in the mixed models. The ER-composition by gender-composition cross-tabulation was summarized descriptively. A Pearson chi-square test was planned only if expected-cell assumptions were adequate.

##### Design Sensitivity and Software

Because we inherited the sample size from the parent study, we evaluated design sensitivity without relying on observed post hoc power. Parametric simulate-refit grids estimated detection rates for standardized average IAPS coefficients in each IPS model. An operational minimum detectable effect required at least 80% all-requested detection for two consecutive positive grid points, at least 95% valid refits, and a beta-zero exact binomial interval containing α =.05. For the small profile groups, we approximated pairwise minimum detectable Hedges’ *g* values from the observed cell sizes and variance ratios. These values describe sensitivity to specified effect sizes and do not establish equivalence when an observed test is nonsignificant.

We conducted all analyses in R (Version 4.4.2). MdRQA analyses were performed using the *crqa* package (2.0.6). We fitted linear mixed-effects models using the *lme4* (1.1-36) and *lmerTest* (3.2-1) packages, applying Satterthwaite denominator degrees of freedom. To compute estimated means, trends, and contrasts, we used *emmeans* (1.10.7). We relied on the *performance* (0.15.3) package for model-performance indices, *nlme* (3.1-166) to fit unequal-residual-variance models, and *rstatix* (0.7.2) for Welch and Games-Howell analyses. We report our results using unstandardized coefficients, standard errors, 95% confidence intervals, test statistics, denominator degrees of freedom, exact *p* values (adjusted where relevant), effect sizes, model *R²*, and ICCs.

## Results

### The Interplay between IPS and IAPS (Hypotheses 1 & 2)

The first aim was to test whether intrapersonal physiological synchrony (IAPS) and interpersonal physiological synchrony (IPS) covary, consistent with synergy rather than a trade-off. To address this question, we assessed the association between IAPS and IPS within each interaction segment and physiological modality.

Initially, same-segment triad-level Pearson correlations (i.e., group-aggregated; Table 1) showed that IAPS was positively associated with IPS-BPM in every segment (average *r* =.52, range =.46–.62); all four correlations remained significant after Holm correction. For IPS-EDA, associations were more context-dependent, with weak or nonsignificant correlations during baseline and the affect manipulation but strong positive associations during the joint task segments (average *r* =.35, range =.14–.61); the T1 and T2 correlations remained significant after Holm correction across the twelve same-segment IAPS–IPS correlations. IPS-RESP was also positively associated with IAPS in all four segments (average *r* =.62, range =.45–.79); all four correlations remained significant after Holm correction, with the largest association during the manipulation (Table 1).

**Table 1.**
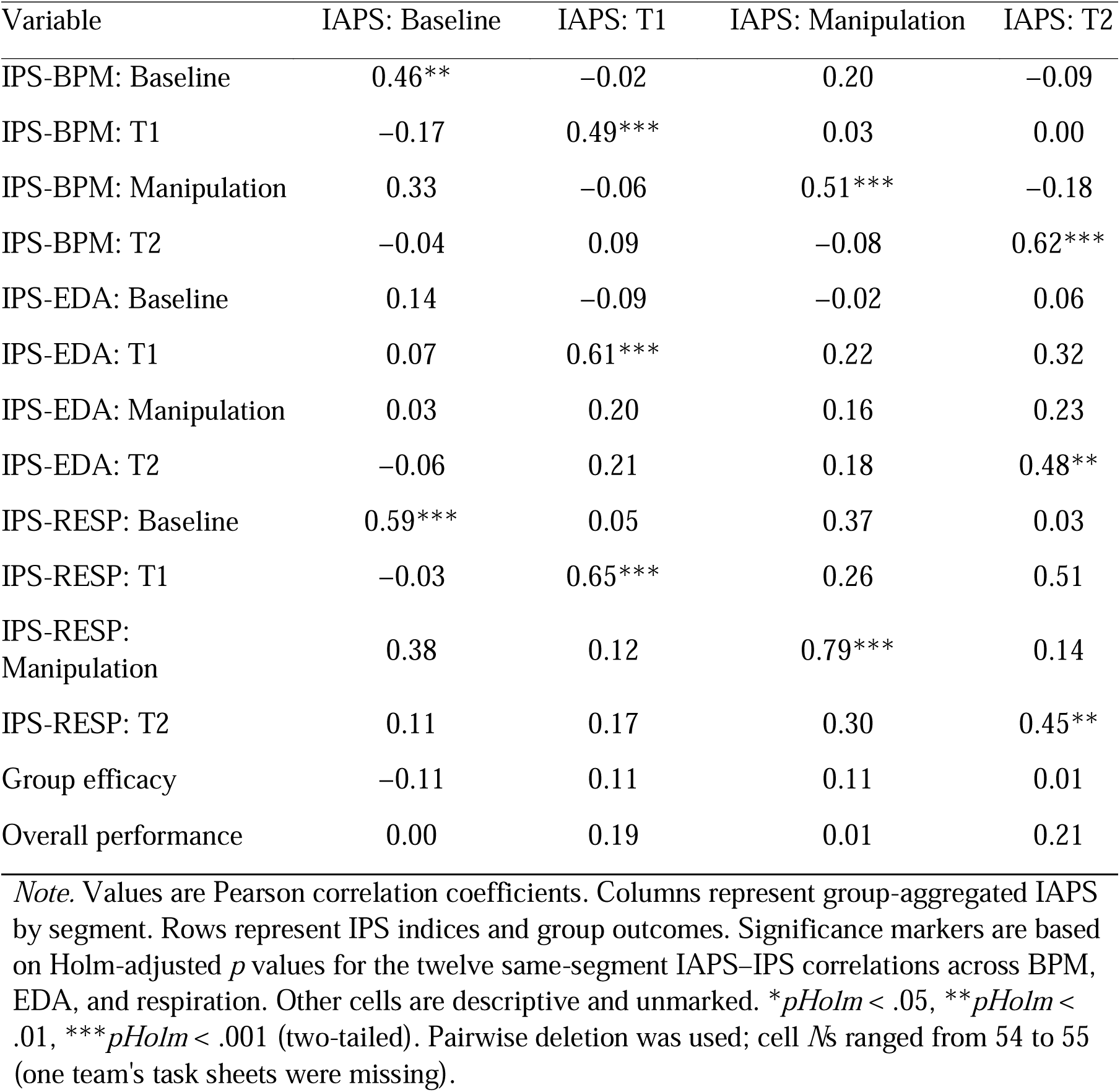
Group-Level Pearson Correlations between IAPS, IPS, and Group Outcomes Across Study Segments.

| Variable | IAPS: Baseline | IAPS: T1 | IAPS: Manipulation | IAPS: T2 |
| --- | --- | --- | --- | --- |
| IPS-BPM: Baseline | 0.46** | −0.02 | 0.20 | −0.09 |
| IPS-BPM: T1 | −0.17 | 0.49*** | 0.03 | 0.00 |
| IPS-BPM: Manipulation | 0.33 | −0.06 | 0.51*** | −0.18 |
| IPS-BPM: T2 | −0.04 | 0.09 | −0.08 | 0.62*** |
| IPS-EDA: Baseline | 0.14 | −0.09 | −0.02 | 0.06 |
| IPS-EDA: T1 | 0.07 | 0.61*** | 0.22 | 0.32 |
| IPS-EDA: Manipulation | 0.03 | 0.20 | 0.16 | 0.23 |
| IPS-EDA: T2 | −0.06 | 0.21 | 0.18 | 0.48** |
| IPS-RESP: Baseline | 0.59*** | 0.05 | 0.37 | 0.03 |
| IPS-RESP: T1 | −0.03 | 0.65*** | 0.26 | 0.51 |
| IPS-RESP: Manipulation | 0.38 | 0.12 | 0.79*** | 0.14 |
| IPS-RESP: T2 | 0.11 | 0.17 | 0.30 | 0.45** |
| Group efficacy | −0.11 | 0.11 | 0.11 | 0.01 |
| Overall performance | 0.00 | 0.19 | 0.01 | 0.21 |

*Note.* Values are Pearson correlation coefficients. Columns represent group-aggregated IAPS by segment. Rows represent IPS indices and group outcomes. Significance markers are based on Holm-adjusted $p$ values for the twelve same-segment IAPS–IPS correlations across BPM, EDA, and respiration. Other cells are descriptive and unmarked. \* $p_{Holm} < .05$ , \*\* $p_{Holm} < .01$ , \*\*\* $p_{Holm} < .001$ (two-tailed). Pairwise deletion was used; cell $N$ s ranged from 54 to 55 (one team's task sheets were missing).

For the primary examinations of Hypotheses 1 and 2, and to model these associations across segments while accounting for repeated observations within triads, we performed mixed-effects models using the outcome-specific random structures described above (55 triads; 220 observations; Table 2; Figure 2). For BPM, IPS-BPM showed a positive average association with group-aggregated IAPS [*b* = 0.58, *SE* = 0.09, 95% CI [0.41, 0.75], *F*(1, 107.12) = 43.91, *p* <.001, partial *R²* =.291, *f²* =.410] and differed by segment [*F*(3, 159.88) = 10.51, *p* <.001]. The IAPS × segment interaction was not significant [*F*(3, 197.26) = 1.97, *p* =.120, partial *R²* =.029, *f²* =.030], indicating that the positive IAPS–IPS-BPM association did not significantly differ across segments. Segment-specific IAPS slopes were positive in all segments with CIs excluding zero (BL: *b* = 0.67, 95% CI [0.38, 0.96]; T1: *b* = 0.68, 95% CI [0.37, 0.99]; MAN: *b* = 0.33, 95% CI [0.07, 0.58]; T2: *b* = 0.64, 95% CI [0.31, 0.97]) and did not differ after Holm correction (smallest *p*holm =.296). The final IPS-BPM model had marginal *R²* =.260, 95% CI [.173,.387], conditional *R²* =.535, 95% CI [.409,.663], and observation-averaged adjusted ICC =.371, 95% CI [.204,.522]. The triad-intercept, within-triad IAPS-slope, and residual standard deviations were 1.39, 0.49, and 2.50, respectively (Table S2).

**Figure 2.**
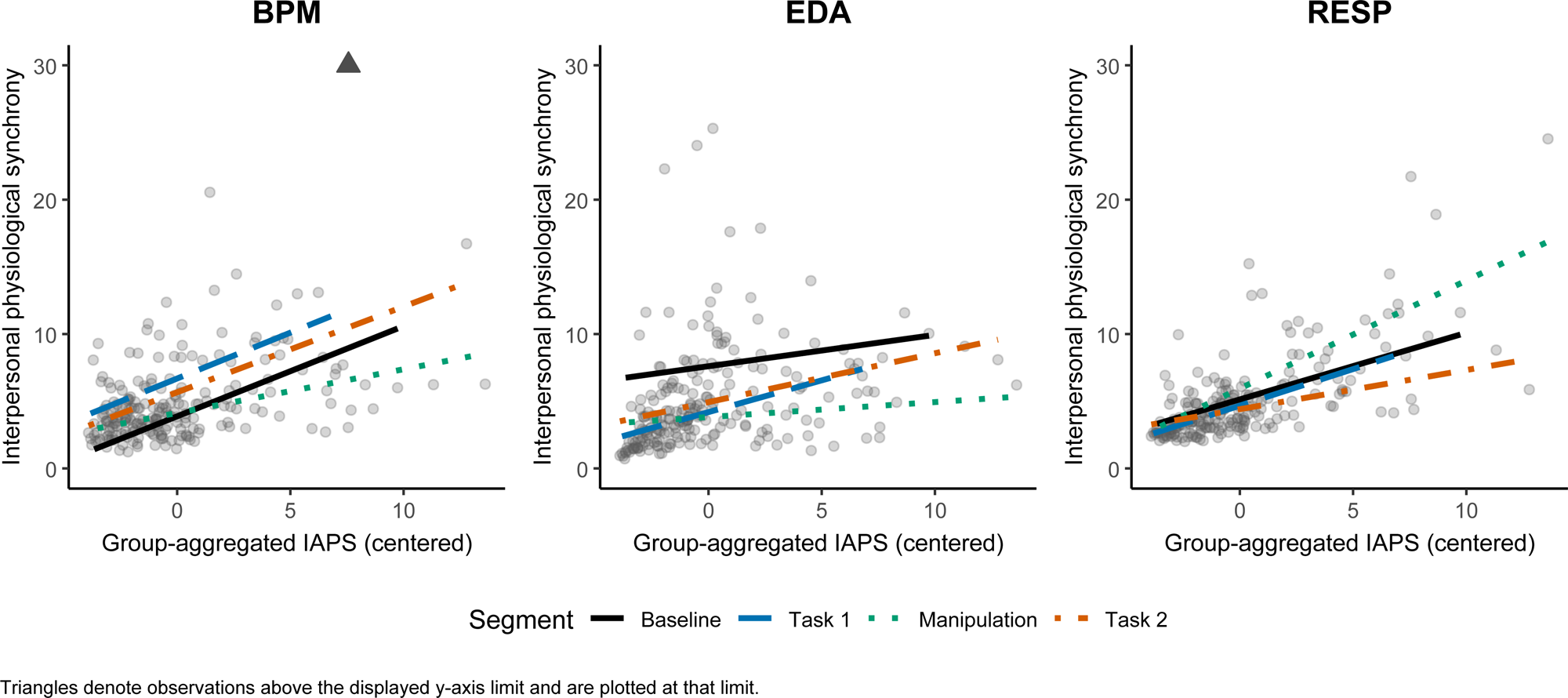
Group-level interpersonal physiological synchrony (IPS) as a function of group-aggregated intrapersonal physiological synchrony (IAPS; centered), shown separately for each interaction segment and physiological modality. Gray points indicate observed group-by-segment values. Colored lines show model-predicted values from the mixed-effects models as a function of centered IAPS, with separate slopes by segment. The y-axis is capped at 30 for comparability across panels; one observation exceeding 30 is shown at the cap with a triangle marker.

**Table 2.** Mixed-effects Models Predicting IPS from Group-Aggregated IAPS Across Interaction Segments.

| Predictor | *b* | *SE* | *df* | *t* | *p* | 95% CI |
| --- | --- | --- | --- | --- | --- | --- |
| IPS-BPM |  |  |  |  |  |  |
| Intercept (grand mean, centered IAPS) | 5.09 | 0.26 | 59.2 | 19.38 | < .001 | [4.56, 5.62] |
| IAPS (average slope; centered) | 0.58 | 0.09 | 107.1 | 6.63 | < .001 | [0.41, 0.75] |
| Segment: BL (sum contrast) | -1.21 | 0.33 | 155.8 | -3.72 | < .001 | [-1.86, -0.57] |
| Segment: T1 (sum contrast) | 1.60 | 0.36 | 169.2 | 4.42 | < .001 | [0.88, 2.32] |
| Segment: MAN (sum contrast) | -0.96 | 0.32 | 154.6 | -2.97 | .003 | [-1.60, -0.32] |
| IAPS × BL (sum contrast) | 0.09 | 0.12 | 203.9 | 0.75 | .451 | [-0.15, 0.33] |
| IAPS × T1 (sum contrast) | 0.10 | 0.12 | 197.4 | 0.83 | .409 | [-0.14, 0.35] |
| IAPS × MAN (sum contrast) | -0.25 | 0.11 | 194.3 | -2.38 | .018 | [-0.46, -0.04] |
| IPS-EDA |  |  |  |  |  |  |
| Intercept (grand mean, centered IAPS) | 5.14 | 0.30 | 59.7 | 17.05 | < .001 | [4.53, 5.74] |
| IAPS (average slope; centered) | 0.30 | 0.07 | 209.9 | 4.05 | < .001 | [0.15, 0.44] |
| Segment: BL (sum contrast) | 2.47 | 0.36 | 161.7 | 6.82 | < .001 | [1.75, 3.18] |
| Segment: T1 (sum contrast) | -0.95 | 0.39 | 168.4 | -2.44 | .016 | [-1.72, -0.18] |
| Segment: MAN (sum contrast) | -1.31 | 0.36 | 164.0 | -3.62 | < .001 | [-2.03, -0.60] |
| IAPS × BL (sum contrast) | -0.06 | 0.12 | 200.4 | -0.52 | .604 | [-0.30, 0.17] |
| IAPS × T1 (sum contrast) | 0.18 | 0.13 | 193.1 | 1.34 | .181 | [-0.08, 0.44] |
| IAPS × MAN (sum contrast) | -0.18 | 0.11 | 185.6 | -1.70 | .090 | [-0.40, 0.03] |
| IPS-RESP |  |  |  |  |  |  |
| Intercept (grand mean, centered IAPS) | 5.04 | 0.17 | 57.9 | 29.34 | < .001 | [4.70, 5.38] |
| IAPS (average slope; centered) | 0.53 | 0.06 | 92.5 | 8.72 | < .001 | [0.41, 0.65] |
| Segment: BL (sum contrast) | 0.10 | 0.23 | 157.9 | 0.41 | .685 | [−0.37, 0.56] |
| Segment: T1 (sum contrast) | −0.37 | 0.26 | 166.0 | −1.40 | .163 | [−0.88, 0.15] |
| Segment: MAN (sum contrast) | 0.90 | 0.23 | 157.5 | 3.91 | < .001 | [0.45, 1.36] |
| IAPS × BL (sum contrast) | −0.04 | 0.09 | 181.7 | −0.42 | .673 | [−0.21, 0.14] |
| IAPS × T1 (sum contrast) | 0.01 | 0.09 | 196.4 | 0.11 | .914 | [−0.17, 0.19] |
| IAPS × MAN (sum contrast) | 0.27 | 0.08 | 186.1 | 3.47 | < .001 | [0.12, 0.42] |

*Note.* Fixed effects are from the selected mixed-effects models ( $N = 55$ triads; 220 triad-segment observations). IPS-BPM used uncorrelated triad random intercepts and within-triad IAPS slopes, IPS-EDA used a triad random intercept, and IPS-RESP used correlated triad random intercepts and within-triad IAPS slopes. IAPS was centered and sum-to-zero contrasts were used for segment. Thus, the intercept and IAPS coefficient represent the grand mean and average IAPS slope across segments. The BL, T1, and MAN coefficients represent deviations from these averages; the corresponding T2 coefficients are implied by the negative sum of the three displayed contrasts.

For EDA, IPS-EDA differed across segments [*F*(3, 163.58) = 16.64, *p* <.001] and showed a positive average association with IAPS [*b* = 0.30, *SE* = 0.07, 95% CI [0.15, 0.44], *F*(1, 209.87) = 16.39, *p* <.001, partial *R²* =.072, *f²* =.078], with no significant segment-related variation in the primary model [*F*(3, 191.56) = 1.28, *p* =.284, partial *R²* =.020, *f²* =.020]. Segment-specific slopes were positive and statistically distinguishable from zero during T1 and T2 (T1: *b* = 0.47, 95% CI [0.16, 0.79]; T2: *b* = 0.36, 95% CI [0.08, 0.65]) but not during baseline or the manipulation (BL: *b* = 0.23, 95% CI [−0.03, 0.50]; MAN: *b* = 0.11, 95% CI [−0.13, 0.35]); none of the six pairwise slope comparisons was significant after Holm correction. The final IPS-EDA model had marginal *R²* =.250, 95% CI [.186,.347], conditional *R²* =.418, 95% CI [.314,.537], and adjusted ICC =.224, 95% CI [.076,.363]. The triad-intercept and residual standard deviations were 1.58 and 2.94, respectively (Table S2).

For respiration, IPS-RESP differed across segments [*F*(3, 160.56) = 6.08, *p* <.001] and showed a positive average association with IAPS [*b* = 0.53, *SE* = 0.06, 95% CI [0.41, 0.65], *F*(1, 92.48) = 76.06, *p* <.001, partial *R²* =.451, *f²* =.822]. The IAPS × segment interaction was significant in the primary model [*F*(3, 183.66) = 4.73, *p* =.003, partial *R²* =.072, *f²* =.077]. Segment-specific slopes were positive in all four segments (BL: *b* = 0.50, 95% CI [0.29, 0.71]; T1: *b* = 0.54, 95% CI [0.32, 0.76]; MAN: *b* = 0.80, 95% CI [0.62, 0.99]; T2: *b* = 0.29, 95% CI [0.05, 0.53]). Only the manipulation slope exceeded the T2 slope after Holm correction (*p* =.003); the other slope comparisons were nonsignificant. The final IPS-RESP model had marginal *R²* =.501, 95% CI [.403,.605], conditional *R²* =.639, 95% CI [.559,.740], and observation-averaged adjusted ICC =.277, 95% CI [.133,.459]. The triad-intercept, within-triad IAPS-slope, and residual standard deviations were 0.85, 0.28, and 1.83, respectively; the intercept-slope correlation was.963 (Table S2).

We further tested whether the IAPS–IPS associations could be attributed to other potentially central factors. The pattern of results was unchanged when adjusting separately for segment-specific mean HR for BPM, tonic skin conductance for EDA, and RSA, triad mean age, pre-task positive and negative affect, and ER composition for all three outcomes (Tables S3–S7). Only tonic skin conductance independently predicted IPS-EDA, *F*(1, 59.55) = 7.96, *p* =.006, yet did not remove the positive IAPS–IPS-EDA association. No covariate had a significant omnibus association with IPS-RESP (all *omnibus p* values ≥.214), and its positive average association with IAPS remained significant in every adjusted model.

All selected primary IPS models converged and were nonsingular (Table S8). Formal residual tests detected nonnormality and unequal residual variance in the raw-scale IPS-BPM, IPS-EDA, and IPS-RESP models (all *p*s <.001; Table S9a; Figures S1–S3). Allowing residual variance to differ by segment improved fit for IPS-BPM (LR = 19.24, *p* <.001), IPS-EDA (LR = 81.21, *p* <.001), and IPS-RESP (LR = 18.79, *p* <.001). The corresponding log1p models no longer showed detectable heteroscedasticity (all *p*s ≥.425), although formal residual-normality tests remained significant (all *p*s ≤.042). The selected RESP random-slope model had a high intercept-slope correlation (*r* =.963), but refits with three optimizers converged, remained nonsingular, and produced practically identical likelihoods, fixed effects, and variance parameters. A targeted uncorrelated random-slope sensitivity retained both the positive average IAPS association and the IAPS × segment interaction (Table S8). Omnibus tests and segment-specific IAPS slopes across IPS model specifications are reported in Tables S9b–S9c.

For IPS-BPM, the positive average IAPS association remained significant in the unequal-variance and log1p models and in all 55 leave-one-triad-out refits (Tables S9b and S10). The within-between decomposition also yielded a positive between-triad effect (*b* = 0.44, 95% CI [0.17, 0.72], *p* =.002) and a positive within-triad effect (*b* = 0.67, 95% CI [0.44, 0.89], *p* <.001; Table S11). The IAPS × segment interaction remained nonsignificant in the unequal-variance (*p* =.104), log1*p* (*p* =.706), and within-between (*p* =.298) models. Across the 55 leave-one-triad-out refits, interaction *p* values ranged from.043 to.173, with 2 refits below.05. Thus, BPM moderation was not supported across model specifications, although this conclusion was not completely invariant to omission of individual triads. The simulation grid reached 87.6% detection at a standardized coefficient of.25, but the beta-zero rejection rate was.080; therefore, no BPM minimum detectable effect met the operational calibration rule (Table S12).

For IPS-EDA, the positive average IAPS association likewise remained significant in the unequal-variance and log1p models and in all 55 leave-one-triad-out refits (Tables S9b and S10). The within-between decomposition yielded a positive between-triad effect (*b* = 0.40, 95% CI [0.09, 0.72], *p* =.014) and a positive within-triad effect (*b* = 0.24, 95% CI [0.08, 0.40], *p* =.003; Table S11). However, the IAPS × segment interaction was significant in the unequal-variance (*p* =.029) and log1*p* (*p* =.021) models but not in the primary raw-scale (*p* =.284) or within-between (*p* =.364) models. Only the T1–MAN slope contrast was significant after Holm correction in the unequal-variance (*p* =.020) and log1*p* (*p* =.013) models (Table S9c). Leave-one-triad-out interaction *p* values ranged from.077 to.396, with no refit below.05. Thus, evidence that the IAPS–IPS-EDA association differed across segments was specification-sensitive. A standardized average IAPS coefficient of.20 was detectable with approximately 90% power (Table S12).

For IPS-RESP, the positive average IAPS association remained significant in the unequal-variance and log1p models and in all 53 valid same-specification leave-one-triad-out refits; two deletion refits were invalid (Tables S9b–S11). The within-between decomposition yielded a positive between-triad effect (*b* = 0.74, 95% CI [0.57, 0.91], *p* <.001) and a positive within-triad effect (*b* = 0.46, 95% CI [0.31, 0.62], *p* <.001). The IAPS × segment interaction was significant in the unequal-variance (*p* <.001) and within-between (*p* =.009) models and in every valid leave-one-triad-out refit (*p* values ranged from <.001 to.028), but not in the log1p model (*p* =.212). All four segment slopes remained positive across specifications, although the unequal-variance T2 slope was not distinguishable from zero. Thus, the average positive IAPS–IPS-RESP association was robust, whereas evidence for segment moderation was specification-sensitive. A standardized average IAPS coefficient of.20 met the operational sensitivity rule with 86.0% all-requested detection, although frequent singular simulated refits make this estimate approximate (Table S12).

Taken together, these findings indicate that intrapersonal and interpersonal synchrony tend to covary. The positive average associations for BPM, EDA, and respiration were robust across the relevant analyses. Segment moderation was not robustly supported for BPM and specification-sensitive for EDA and respiration.

### Contextual Dynamics and Individual Differences in IAPS (Hypothesis 3)

To examine Hypothesis 3, i.e., whether IAPS is stable across contexts or varies with interaction demands, we tested segment differences and stability across the study segments. We evaluated (a) associations between IAPS levels across interaction segments, (b) mean differences in IAPS levels across segments, and (c) associations between IAPS and individual-difference measures. Observed means (*SD*s) were BL = 5.91 (5.63), T1 = 3.55 (4.14), MAN = 6.03 (5.84), and T2 = 4.51 (4.93).

IAPS levels during the baseline measurement and the negative affect induction manipulation were significantly and positively correlated, *r* =.34, 95% CI [.20,.47], *pHolm* <.001, as were IAPS levels during both task trials, *r* =.40, 95% CI [.26,.52], *pHolm* <.001. The remaining correlations were nonsignificant (Table 3).

**Table 3.** Individual-Level Pearson Correlations of IAPS Across Segments and With Individual-Difference Self-Report Measures.

| Variable/Segment | IAPS:<br>Baseline | IAPS: T1 | IAPS:<br>Manipulation | IAPS: T2 |
| --- | --- | --- | --- | --- |
| IAPS: Baseline | - | −0.06 | 0.34*** | −0.01 |
| IAPS: T1 | −0.06 | - | 0.10 | 0.40*** |
| IAPS: Manipulation | 0.34*** | 0.10 | - | 0.14 |
| IAPS: T2 | -0.01 | 0.40*** | 0.14 | - |
| Big Five:<br>Extraversion | 0.01 | -0.15 | 0.15 | -0.09 |
| Big Five:<br>Agreeableness | -0.07 | -0.09 | -0.02 | -0.13 |
| Big Five:<br>Conscientiousness | -0.03 | 0.05 | 0.02 | 0.11 |
| Big Five: Neuroticism | 0.08 | 0.00 | -0.05 | -0.11 |
| Big Five: Openness | 0.09 | -0.03 | 0.09 | -0.07 |
| Emotion regulation<br>difficulties (DERS) | -0.03 | 0.10 | -0.12 | 0.03 |
| Social anxiety | -0.02 | 0.03 | -0.10 | -0.09 |
| Generalized anxiety | 0.01 | 0.05 | -0.04 | 0.04 |
| Depressive symptoms | 0.08 | 0.06 | -0.12 | -0.06 |
| Life satisfaction | -0.02 | -0.10 | -0.04 | -0.04 |
| Emotion regulation<br>(ERQ): Cognitive<br>reappraisal | -0.03 | 0.00 | 0.03 | 0.03 |
| Emotion regulation<br>(ERQ): Emotion<br>suppression | -0.01 | -0.03 | -0.17 | -0.01 |
| Negative mood<br>regulation | 0.00 | -0.07 | 0.08 | -0.08 |
| Differentiation of<br>Self: Emotional<br>reactivity | -0.07 | 0.10 | 0.08 | 0.09 |
| Differentiation of<br>Self: I-position | -0.03 | 0.15 | 0.02 | 0.08 |
| Differentiation of<br>Self: Emotional cutoff | 0.05 | 0.14 | 0.03 | 0.03 |
| Differentiation of<br>Self: Fusion with<br>others | -0.03 | 0.15 | 0.10 | 0.10 |
| Emotion contagion | -0.16 | 0.05 | -0.03 | -0.04 |
| Attachment anxiety | 0.04 | 0.10 | −0.05 | 0.09 |
| Attachment avoidance | 0.02 | −0.07 | −0.07 | −0.01 |
| Self-deceptive<br>enhancement | −0.11 | 0.00 | −0.04 | −0.03 |
| Impression<br>management | 0.12 | 0.01 | 0.01 | 0.02 |

*Note.* Values are Pearson correlation coefficients. The six cross-segment IAPS correlations were Holm-adjusted as one stability family. The 88 correlations between segment-specific IAPS and individual-difference measures were controlled using the Benjamini–Hochberg false-discovery-rate procedure. Significance markers reflect adjusted $p$ values; no individual-difference correlation survived false-discovery-rate correction. \* $p_{adjusted} < .05$ , \*\* $p_{adjusted} < .01$ , \*\*\* $p_{adjusted} < .001$ (two-tailed). Cell $N$ s ranged from 161 to 165 because of questionnaire missingness.

We then tested segment effects using a mixed-effects model with random intercepts for triad and participant nested within triad (55 triads; 165 participants; 660 person-segment observations). IAPS levels differed significantly by segment [*F*(3, 492.00) = 10.28, *p* <.001, partial *R²* =.059, *f²* =.063]. Holm-corrected contrasts indicated that IAPS levels during BL were higher compared to T1 (Δ = 2.37, *p* <.001) and T2 (Δ = 1.40, *p* =.024), and MAN exceeded T1 (Δ = 2.49, *p* <.001) and T2 (Δ = 1.52, *p* =.016), whereas BL and MAN did not differ (Δ = 0.12, *p* =.820) and T1 and T2 did not differ (Δ = 0.97, *p* =.133) (Figure 3). Thus, IAPS levels were related within similar contexts and were higher during passive co-presence than during active joint tasks.

**Figure 3.**
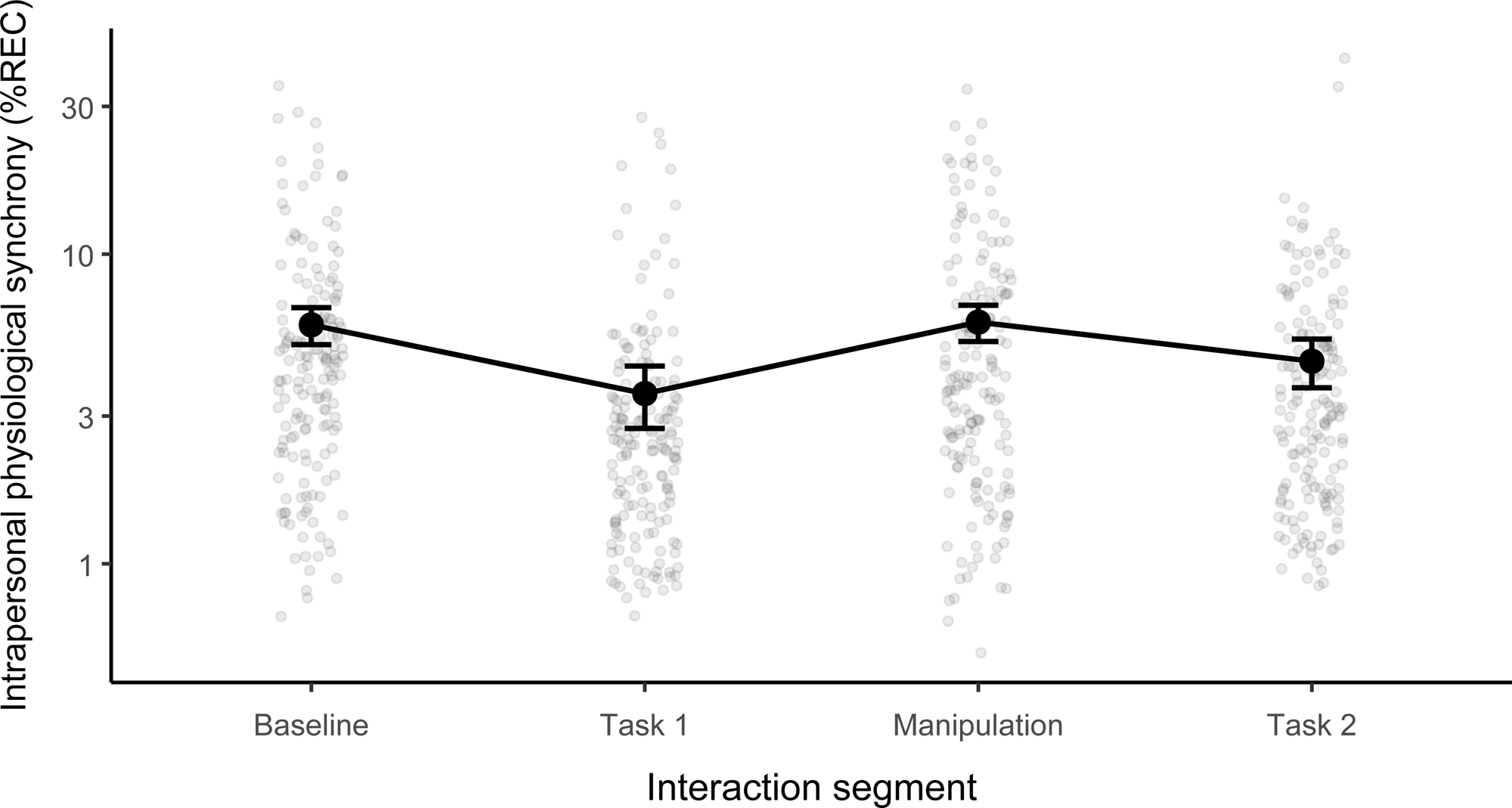
Within-person differences in IAPS levels across segments. Points represent individual IAPS observations. Solid points connected by a line represent estimated marginalmeans from the mixed-effects model, and error bars represent 95% confidence intervals. The y-axis is log10-scaled for display; the estimated marginal means and confidence intervals were obtained from the primary raw-scale model.

The selected participant-level IAPS model converged and was nonsingular (Table S8). Formal residual tests detected nonnormality and unequal residual variance in the raw-scale model (both *p*s <.001; Table S9a; Figures S1–S3). Allowing residual variance to differ by segment improved fit (LR = 23.93, *p* <.001). The log1p model no longer showed detectable heteroscedasticity (*p* =.274), although formal residual nonnormality remained significant (*p* <.001). Omnibus segment tests across specifications are reported in Table S9b.

The final segment model had marginal *R²* =.038, 95% CI [.018,.072], conditional *R²* =.183, 95% CI [.107,.265], and adjusted ICC =.151, 95% CI [.073,.232]. The triad-intercept, participant-intercept, and residual standard deviations were 0.71, 1.88, and 4.77, respectively (Table S2). The omnibus segment effect remained supported in the unequal-variance, log1p, and leave-one-triad-out analyses. Higher IAPS levels in passive than active segments remained supported in the unequal-variance and log1p models and were largely retained in leave-one-triad-out analyses: three passive–active contrasts remained Holm-significant in all 55 refits, whereas BL–T2 did so in 51. The T1–T2 contrast was significant only in the log1p model (Tables S9b and S10).

We then tested the associations between IAPS levels throughout the study’s segments and the Big Five personality traits, emotion regulation (ER) abilities, social inclinations (attachment, emotion contagion, differentiation of self), and psychopathology. IAPS levels during baseline and the manipulation were negatively correlated with reported emotion contagion, *r* = −.16, *p* =.036, and emotion suppression, *r* = −.17, *p* =.030, respectively. However, neither association survived Benjamini–Hochberg correction (both *pBH* =.845), and no other individual-difference correlation was significant after correction (Table 3).

Lastly, we compared IAPS levels between participants below versus at or above the established cutoffs for significant difficulties in ER and probable common affective disorders (i.e., social anxiety, generalized anxiety, and depression; Welch independent-samples *t* tests, two-sided; Table S13). None of the 16 comparisons was significant after Holm correction. Participants with probable depression had nominally lower IAPS levels during T2 [*t*(157.94) = −2.19, *p* =.030, *Hedges’ g* = −0.32], but this difference did not survive correction, *pHolm* =.478.

Taken together, the findings presented thus far suggest that IAPS varies depending on the context of the group interaction, with complex state-trait characteristics. Participants’ relative IAPS levels were stable across similar contexts, and mean IAPS was higher during passive co-presence than during active joint tasks. However, IAPS was not reliably associated with the broader individual-difference measures or clinical-threshold classifications after correction for multiple testing.

Because triads were intentionally formed under three ER-composition conditions, ER composition was examined as a secondary predictor of IAPS. ER composition had an omnibus association with IAPS averaged across segments [*F*(2, 52.00) = 3.32, *p* =.044, partial *R²* =.113, *f²* =.128]. Estimated marginal means were 4.68, 5.98, and 4.52 for compositions with zero, one, and two members with self-reported elevated ER difficulties, respectively, but none of the three pairwise comparisons was significant after Holm correction (all *ps* ≥.061). The ER-composition × segment interaction was also nonsignificant, *F*(6, 486.00) = 1.70, *p* =.119 (Table S14).

### How do IAPS and IPS relate to Group Functioning? (Hypothesis 4)

Finally, we explored whether different combinations of intrapersonal and interpersonal synchrony during task performance were associated with group outcomes. For this exploratory analysis, IAPS and IPS were averaged across the two task segments and dichotomized at the median to form high and low classifications. Groups were then categorized into four combinations: high IAPS–high IPS (HH), high IAPS–low IPS (HL), low IAPS–high IPS (LH), and low IAPS–low IPS (LL). Overall, 63.6% of the groups (*n* = 35) were classified into the same category in both composite variables (i.e., IAPS × IPS-BPM and IAPS × IPS-EDA), predominantly the HH (29.1%) and LL (25.5%) subgroups. Objective performance comparisons used 54 complete cases, efficacy comparisons used all 55 triads.

We then performed four Welch one-way ANOVAs (two profile variables × two dependent variables: efficacy and overall task performance; homogeneity of variances not assumed) to explore associations between the high-low profiles and group outcomes. Holm correction was applied across the four omnibus tests, and Games–Howell comparisons were interpreted only when both the corresponding omnibus test remained significant after Holm correction and the Games–Howell-adjusted pairwise *p* value was below.05. Complete findings are reported in Tables S15a–S15c and visualized in Figure 4.

**Figure 4.**
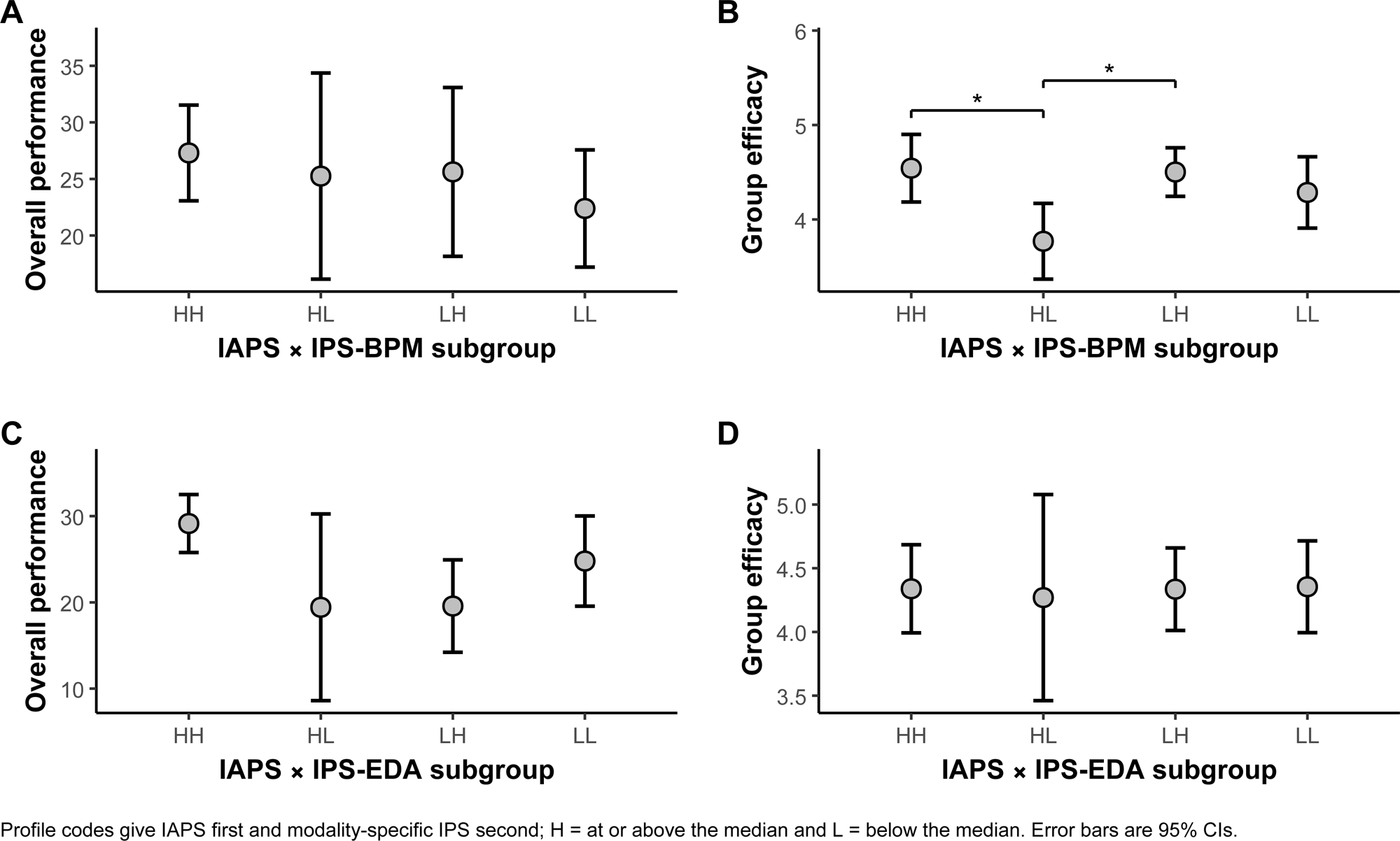
Group outcomes as a function of high–low classifications of intrapersonal physiological synchrony (IAPS) and interpersonal physiological synchrony (IPS). Groups were categorized into four subgroups: HH (high IAPS/high IPS), HL (high IAPS/low IPS), LH (low IAPS/high IPS), and LL (low IAPS/low IPS). Panels show (A) overall task performance by IAPS × IPS-BPM subgroup, (B) group efficacy by IAPS × IPS-BPM subgroup, (C) overall task performance by IAPS × IPS-EDA subgroup, and (D) group efficacy by IAPS × IPS-EDA subgroup. Points represent subgroup means and error bars represent 95% confidence intervals. Asterisks are shown only for Games–Howell-adjusted pairwise comparisons with *p* <.05 when the corresponding Welch omnibus test also remained significant after Holm correction. Welch one-way ANOVAs were used because of heterogeneous variances and unequal subgroup sizes.

Overall performance differed among the IAPS × IPS-EDA profiles before, but not after, Holm correction [*F*(3, 17.42) = 4.43, *p* =.017, *pHolm* =.052, descriptive η² =.156]. Accordingly, the nominal HH–LH performance comparison was not interpreted as a corrected finding. No group differences were observed for group efficacy [*F*(3, 19.38) = 0.02, *p* =.997, *pHolm* = 1.000, descriptive η² =.001].

For the IAPS × IPS-BPM profiles, group efficacy differed after Holm correction [*F*(3, 23.88) = 4.75, *p* =.010, *pHolm* =.039, descriptive η² =.131]. Specifically, the HL subgroup reported lower efficacy than the HH subgroup [HH–HL mean difference = 0.77, 95% CI [0.10, 1.45], *p* =.021, *Hedges’ g* = 1.18] and the LH subgroup [LH–HL mean difference = 0.73, 95% CI [0.13, 1.33], *p* =.016, *Hedges’ g* = 1.72]. Overall performance did not differ across the IAPS × IPS-BPM profiles [*F*(3, 19.49) = 0.75, *p* =.538, *pHolm* = 1.000, descriptive η² =.046].

The primary BPM/EDA profile analyses had limited design sensitivity because subgroup sizes ranged from 7 to 21. At an unadjusted α =.05, the approximate pairwise minimum detectable absolute Hedges’ *g* values ranged from 0.88 to 1.61. Using α =.05/6 as a conservative six-comparison benchmark, they ranged from 1.12 to 2.25. Nonsignificant profile comparisons therefore should not be interpreted as evidence of equivalence (Table S16).

## Discussion

As we work in teams, our physiology is affected both by our internal processes and resource allocation and by our tendency to synchronize with our teammates to facilitate bonding, understanding, and task success. This alignment of physiological operations, referred to as interpersonal physiological synchrony (IPS), has been extensively studied. However, the role of intrapersonal physiological synchrony (IAPS), within-person temporal coordination of activities of both the parasympathetic and sympathetic branches of the autonomic nervous system, has been underexplored. In the current study, we first tested whether IPS and IAPS can coexist. That is, can people’s physiology be both coordinated at an individual level and at the same time synchronized with that of their teammates? We aimed to answer whether there is a ‘trade-off’ (i.e., competition) between the two, or ‘synergy’ (i.e., complementarity). Second, since there is limited understanding of how subsystems of the autonomic nervous system within individuals are impacted by varying social demands of the context, we tested IAPS context-dependency and trait-like characteristics. Lastly, we explored whether a specific combination of IPS and IAPS (e.g., being high on both) is associated with better team outcomes, namely higher subjective team efficacy and objective task performance.

Across segments on average, IAPS was positively associated with IPS in heart rate, EDA, and respiration. These findings support a synergistic complementary relationship in which higher internal autonomic synchrony occurs simultaneously with higher IPS. In other words, there is a synergy, not a trade-off, between these processes. Thus, these findings are consistent with perspectives positing that coordination among autonomic subsystems supports adaptive regulation and social engagement (e.g., Thayer & Lane, 2000). Importantly, the observed associations indicate that IAPS and IPS are related but nonredundant processes. The within-between decomposition reinforced the synergy interpretation by showing positive IAPS–IPS associations both between triads and within the same triad across segments, reducing the possibility that the overall association reflected only stable between-triad differences.

Notwithstanding, the dynamics of IAPS and IPS, and their relationship, presented more complex dynamics. Specifically, IAPS levels varied depending on whether participants were engaged in active joint tasks or passive shared experiences. While associations between IAPS and heart rate synchrony (IPS-BPM) remained consistently positive, the primary segment-specific estimates for sympathetic synchrony (IPS-EDA) were significant during active teamwork but not during the passive segments. This pattern raises the possibility of a “task-orientation” effect, i.e., that IAPS may have several functions. One may be facilitating adaptive social engagement and team socio-emotional connectivity, which may occur to some degree in every interaction and may therefore help explain the cross-setting association with heart rate synchrony, a form of coordination associated with social cohesion (Palumbo et al., 2017). Another possible function is task-related, such that alignment between internal regulatory states and sympathetic IPS dynamics may become more pronounced under conditions of a shared goal toward which attention and arousal are directed. This pattern suggests that the coupling between IAPS and sympathetic IPS, often associated with arousal, attention, and task engagement, may vary depending on interaction and task demands. The respiratory analysis showed another pattern: IAPS–IPS-RESP slopes were positive in every segment and strongest during the manipulation, but the omnibus interaction was not significant after log transformation.

However, evidence that the IAPS–IPS association varied across contexts differed by physiological modality and model specification: differences among the EDA slopes were not significant in the primary model, and the omnibus segment interaction was significant only in the unequal-variance and log1p sensitivity models. The task-orientation interpretation therefore remains provisional and is consistent with broader evidence that coordination dynamics depend on task demands and interaction structure (Gordon et al., 2025).

Beyond showing that IAPS can co-occur with IPS, the findings position IAPS as a unique, context-sensitive phenomenon with both state-and trait-like features, providing theoretical and methodological opportunities. IAPS showed consistency within similar contexts: IAPS measured during active segments correlated with other active segments, and likewise for passive segments, but not across contexts. In addition, IAPS levels during passive interaction segments were higher compared to during task performance. Together, these findings indicate that intrapersonal autonomic coordination is not solely a fixed individual trait, but also a context-sensitive mode of regulation that may reorganize in response to situational demands.

Consistent with this interpretation, IAPS was not reliably associated with the broader individual-difference measures after multiplicity correction. These null associations should not, however, be interpreted as evidence that trait-like influences are absent. They may also have been affected by the recruitment design, which selected participants partly on the basis of ER scores and therefore altered the distribution of ER difficulties relative to the screened population. ER composition had an omnibus association with IAPS averaged across segments, although none of the pairwise contrasts was significant and neither was the ER-composition × segment interaction. Finally, the positive IAPS–IPS associations remained when recorded ER composition was included as an additive covariate.

Reduced IAPS during active coordination should not be interpreted as a maladaptive breakdown of regulation. Instead, it may reflect an adaptive response to increased cognitive and attentional demands and the need for ongoing coordination with teammates (Sebanz et al., 2006). These demands may make flexibility more adaptive than tightly coupled internal dynamics, as effective coordination often requires role differentiation, independent monitoring, and rapid adjustment, potentially resulting in a temporary loosening of intrapersonal synchrony. In contrast, during passive shared-attention contexts such as film viewing, regulatory demands may be comparatively low and there is little functional pressure to segregate from others. Under such conditions, individuals can remain oriented toward the same external stimulus with minimal need for role differentiation, potentially allowing higher internal autonomic coordination to co-occur with interpersonal synchrony (Gordon et al., 2025). From dynamical systems (Kelso et al., 2013) and allostatic (McEwen & Wingfield, 2003) approaches and contemporary work on intrapersonal and interpersonal emotion regulation (Aldao et al., 2015; Chen et al., 2024; Petrova & Gross, 2023; Pruessner et al., 2026), adaptive regulation is characterized less by uniformity than by flexible reconfiguration in response to situational demands. The observed attenuation of IAPS during active interaction may therefore be more consistent with functional reorganization than with regulatory depletion. This interpretation is also consistent with emerging network-physiology perspectives, which suggest that physiological states are characterized by specific patterns of coupling between subsystems (Ivanov, 2021; Ivanov et al., 2016). Together, these findings highlight the value of modeling intrapersonal and interpersonal synchrony across contexts as a multilevel regulatory process rather than a single interpersonal phenomenon.

The MdRQA outcome also warrants a bounded interpretation. %REC indexes how often the reconstructed multivariate system revisits similar states and therefore reflects system-level recurrence or temporal regularity, rather than a process-specific measure of regulation. Importantly, MdRQA recurrence does not require the component signals to match one another at a given time; recurring configurations may include both synchronous and asynchronous relations among the component systems (Amon et al., 2019). We therefore use synchrony as an operational description and do not infer a specific regulatory mechanism from %REC alone.

In the exploratory profile analyses, only the omnibus difference in group efficacy across the IAPS × IPS-BPM profiles survived Holm correction. Both the HH and LH groups reported higher efficacy than the HL group, indicating that this finding was not specific to the HH profile. The EDA-based performance difference did not survive omnibus correction. Given the small and unequal profile groups and their limited design sensitivity, these findings are preliminary and do not establish that concurrently high IAPS and IPS produce better outcomes. These findings contribute to ongoing efforts to identify biobehavioral mechanisms that distinguish effective from less effective groups (Gordon, 2025; Mayo et al., 2021). Further work should examine whether these preliminary profile differences replicate in cooperative contexts that involve shared goals and moment-to-moment coordination demands alongside differentiated problem-solving and independent thinking.

### Limitations

Several limitations should be noted. First, because the design was correlational, we could not infer causality in the IAPS-IPS link. It remains unclear whether stronger IAPS facilitates IPS, whether successful group synchronization stabilizes internal dynamics, or whether both are influenced by additional factors. Experimental designs or longitudinal approaches aimed at generating, controlling, or modeling temporal relations between interpersonal synchrony (Basile et al., 2022; Cirelli et al., 2016) and IAPS could help clarify these relationships. Future studies could move beyond the present correlational approach by modeling self-and co-regulatory change simultaneously, for example with multivariate latent change score models that distinguish multiple coordination patterns over time (Butner et al., 2014).

Second, in the IAPS-IPS analyses, IAPS was aggregated at the group level, which limited our ability to examine within-group heterogeneity. As recent work on individuals’ distinctive ‘synchrony signatures’ suggests (Zilcha-Mano, 2025; Zilcha-Mano et al., 2025), future research should examine whether particular members disproportionately shape collective dynamics and whether their IAPS plays a disproportionate role.

Third, the sample size and distribution of groups in each IAPS-IPS subdivision (e.g., high IAPS and high IPS) were not sufficient to allow a robust comparison between the four combinations with regard to team outcomes.

Fourth, the negative affect induction limits the scope of inference. The study included no positive, neutral, or arousal-matched control condition. Because EDA is sensitive to sympathetic arousal (Dawson et al., 2017), effects involving the manipulation cannot be attributed specifically to negative valence. The film may also have elicited self-referential or perseverative cognition that carried into T2. The findings therefore should not be assumed to generalize to positive affect or to isolate emotional valence from arousal and carryover.

Fifth, the sample was predominantly female (77%). The 55 triads included 2 with no women, 8 with one woman, 16 with two women, and 29 with three women. Triads were formed according to the parent study’s recorded ER-composition assignment, without deliberate gender matching or balancing. Gender composition was not modeled, so its possible influence on IPS and group dynamics cannot be evaluated. Recruitment also targeted specified ER ranges. Although ER composition was examined statistically, these design features limit conclusions about IAPS as a state-versus trait-like construct.

Finally, the sample consisted primarily of healthy young adults with no close ties to one another, limiting the external validity of our findings. Examining clinical populations, long-term working teams in their natural environments and other joint action contexts may reveal circumstances under which intrapersonal and interpersonal synchrony decouple, with implications for social functioning, team success, and mental health.

### Conclusions

In conclusion, this study suggests that understanding group dynamics in general, and interpersonal physiological synchrony in particular, may benefit from considering the internal organization of the autonomic system at the individual level. Rather than competing processes, intrapersonal and interpersonal synchrony were positively associated and are consistent with complementary patterns of physiological organization that vary in response to interaction demands. Effective coordination, particularly in team goal-directed tasks, may be associated with the regulatory flexibility to maintain sufficient internal autonomic organization while adapting to when and how synchronization with teammates is most beneficial. Finally, conceptualizing IAPS as a state–trait phenomenon offers a useful framework for deciphering the complex intrapersonal, interpersonal, and contextual determinants of physiological coordination and their outcomes in social interaction.

## Author Contribution

C.E.: Conceptualization, Methodology, Formal Analysis, Investigation, Visualization, Writing– original draft.

S.W.: Conceptualization, Methodology, Supervision, Software, Writing–review & editing.

I.G.: Conceptualization, Methodology, Supervision, Funding acquisition, Writing–review & editing.

## Data Availability Statement

The data and code supporting the findings of this study are available at https://osf.io/6zwmp/.

## Funding

Ilanit Gordon was funded by ISF grants #434/21.

## Conflicts of Interest

The authors declare no conflicts of interest.

## Notes

https://osf.io/6zwmp/

### Competing Interest Statement

The authors have declared no competing interest.

### Summary of Updates

The manuscript has been revised mainly to clarify the concept and measurement of intrapersonal physiological synchrony, strengthen the statistical analysis and reporting, and add a parallel analysis of interpersonal respiratory synchrony.

