## Supplementary_Materials_r for "There is an “I” in team: Intrapersonal physiological synchrony complements interpersonal physiological synchrony in team interaction"

**MdRQA Parameter Selection**

The parameter-selection table below documents the workflow used before the inferential analyses. For IAPS, each multidimensional matrix contained one participant's BPM, respiration, and EDA. For IPS-BPM, IPS-EDA, and IPS-RESP, each matrix contained the same modality from the three members of a triad. The crqa::mdDelay and crqa::mdFnn functions were used, so both the AMI-based delay estimates and the FNN curves were multidimensional.

Delay was estimated for every analytic unit within each segment, using the first local AMI minimum as the criterion. The same multidimensional procedure was used for IAPS and all three IPS modalities. Segment-level distributions and means were then compared before one common delay was selected for each analysis. The four IAPS means were 8.64, 7.22, 11.53, and 7.21 (mean across segment means = 8.65; selected *d* = 8 as best representation). The corresponding IPS-BPM means were 7.36, 7.80, 8.58, and 7.84 (mean = 7.90; selected *d* = 8); the IPS-EDA means were 15.62, 10.69, 23.20, and 10.58 (mean = 15.02; selected *d* = 15); and the IPS-RESP means were 2.93, 2.98, 2.87, and 3.00 (mean = 2.95; selected *d* = 3). FNN curves were inspected across analytic units and segments, yielding *m* = 10 for IAPS, *m* = 8 for IPS-BPM, *m* = 3 for IPS-EDA, and *m* = 2 for IPS-RESP.

A single parameter set was applied to all units and segments within an analysis. This choice improved comparability, because between-unit or between-segment differences in %REC could not arise from different embedding parameters, but it did not optimize the reconstruction separately for every participant or triad. Radius was tuned manually and iteratively until the average of the four segment-level mean %REC values was approximately 5%. Individual %REC values were therefore free to vary and were used as the analytic outcome. The plots and calculations below show the evidence used for these decisions.

| **MdRQA Parameter-Selection Diagnostics** | | | | |
| --- | --- | --- | --- | --- |
|  | IAPS | | | |
|  | BL | T1 | MAN | T2 |
| Delays | 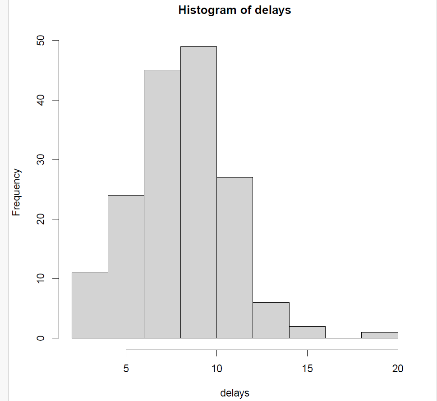 | 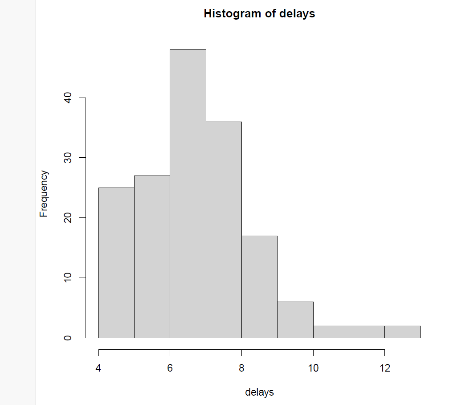 | 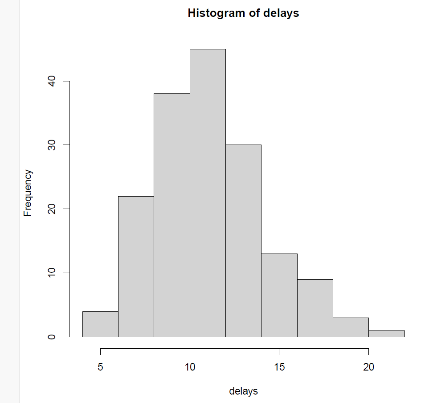 | 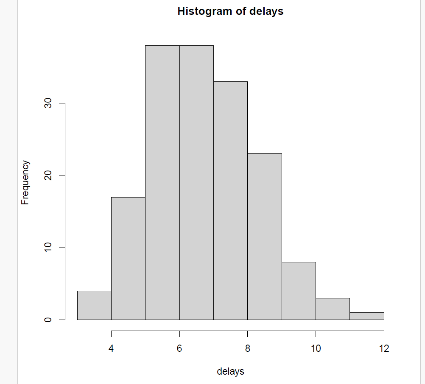 |
|  | Max = 20, mean = 8.636364, *SD* = 2.67119 | Max = 13, mean = 7.224242, *SD* = 1.657766 | Max = 22, mean = 11.52727, *SD* = 3.069447 | Max = 12, mean = 7.206061, *SD* = 1.598387 |
|  | Mean across segment means = 8.648484; **selected *d* = 8** | | | |
| False nearest neighbors | 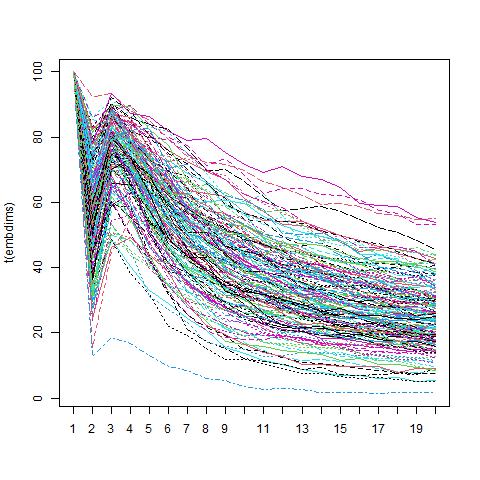 | 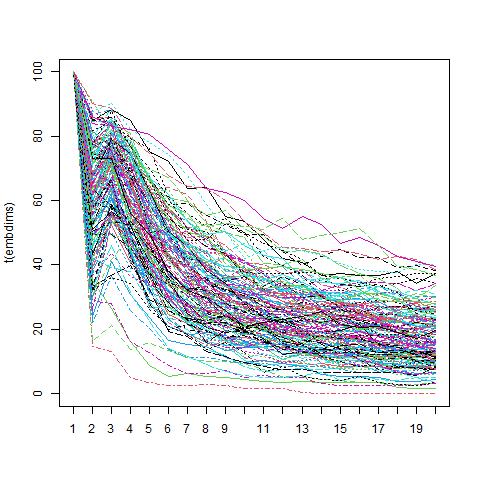 | 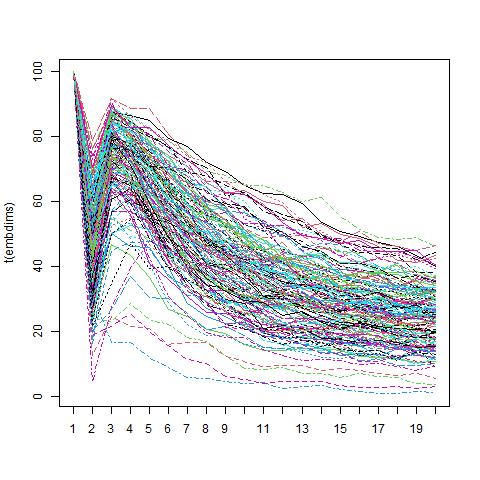 | 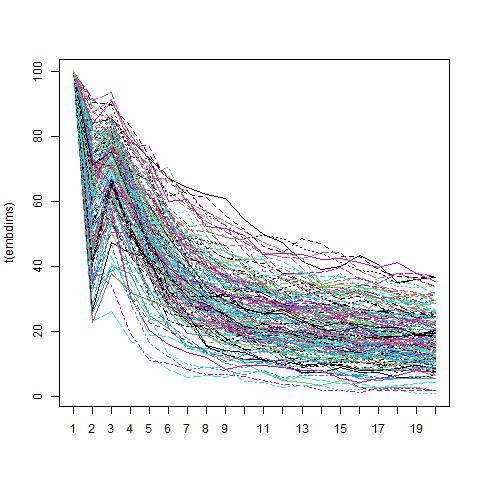 |
|  | **Selected *m* = 10** | | | |
| Radius | With *d* = 8 and *m* = 10, the selected radius was *r* = 4.457. The segment means were 5.912707, 3.545203, 6.032106, and 4.511657; their average was 5.000418% REC. | | | |

|  | IPS-BPM | | | |
| --- | --- | --- | --- | --- |
|  | BL | T1 | MAN | T2 |
| Delays | 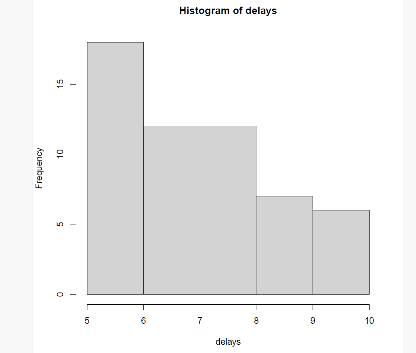 | 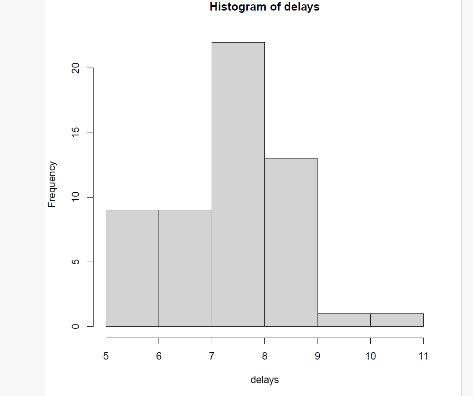 | 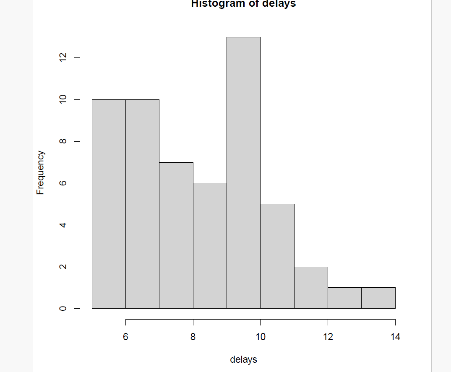 | 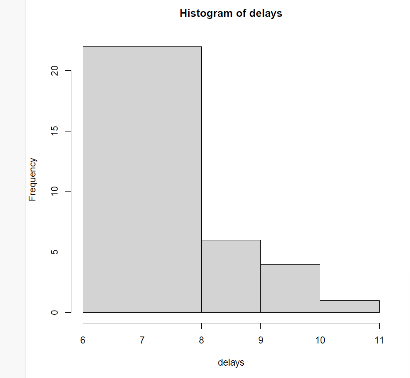 |
|  | Max = 10, mean = 7.363636, *SD* = 1.507557 | Max = 11, mean = 7.8, *SD* = 1.207998 | Max = 14, mean = 8.581818, *SD* = 2.140361 | Max = 11, mean = 7.836364, *SD* = 1.084634 |
|  | Mean across segment means = 7.895455**; selected *d* = 8** | | | |
| False nearest neighbors | 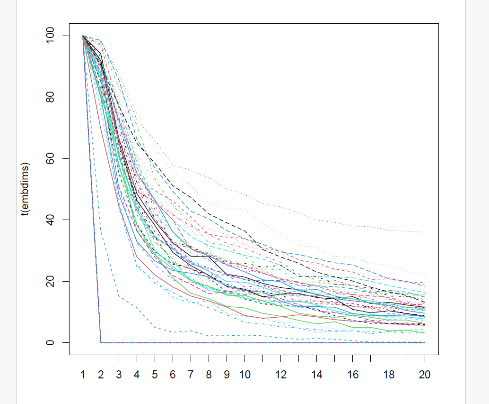 | 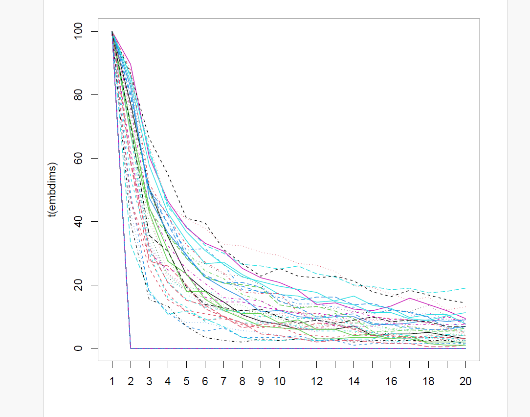 | 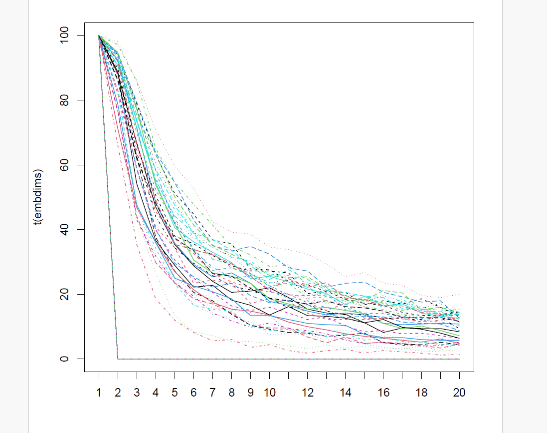 | 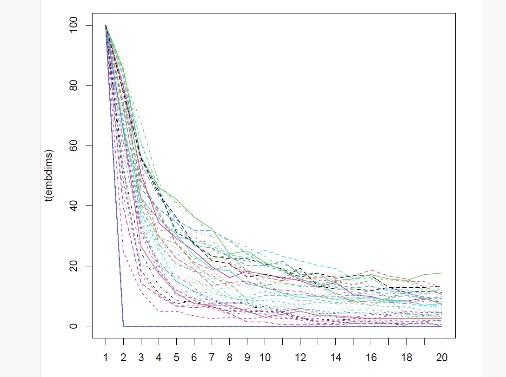 |
|  | **Selected *m* = 8** | | | |
| Radius | With *d* = 8 and *m* = 8, the selected radius was *r* = 4.441. The segment means were 4.503638, 5.555257, 4.679434, and 5.265867; their average was 5.001049% REC. | | | |

|  | IPS-EDA | | | |
| --- | --- | --- | --- | --- |
|  | BL | T1 | MAN | T2 |
| Delays | 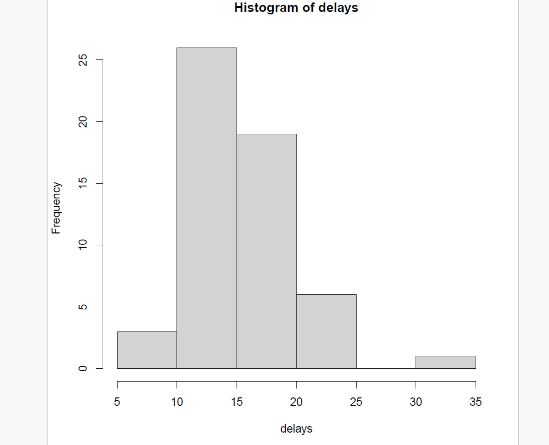 | 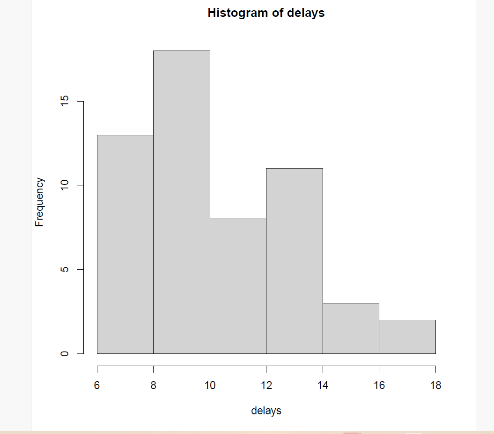 | 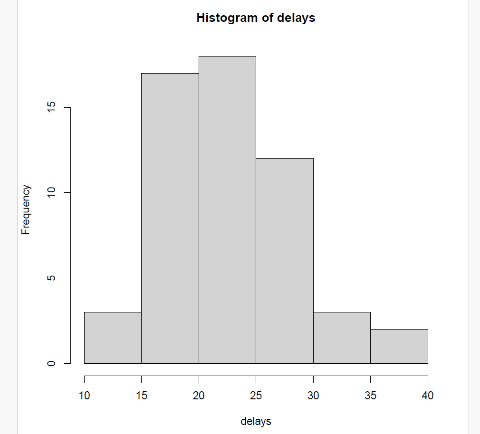 | 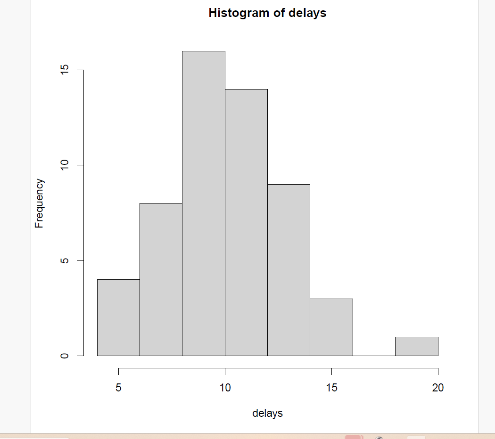 |
|  | Max = 33, mean = 15.61818, *SD* = 4.309726 | Max = 18, mean = 10.69091, *SD* = 2.85367 | Max = 40, mean = 23.2, *SD* = 5.664706 | Max = 19, mean = 10.58182, *SD* = 2.766807 |
|  | Mean across segment means = 15.02273; **selected *d* = 15** | | | |
| False nearest neighbors | 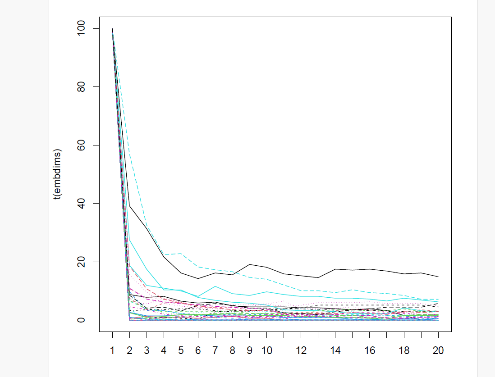 | 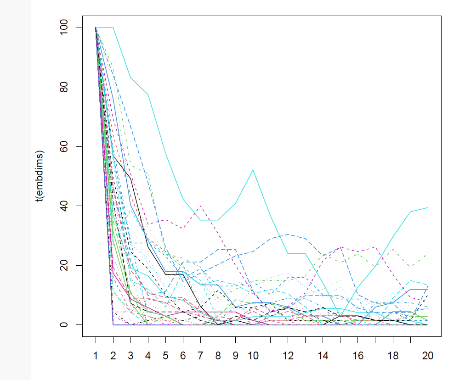 | 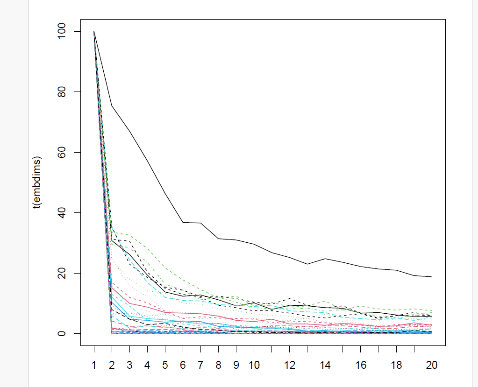 | 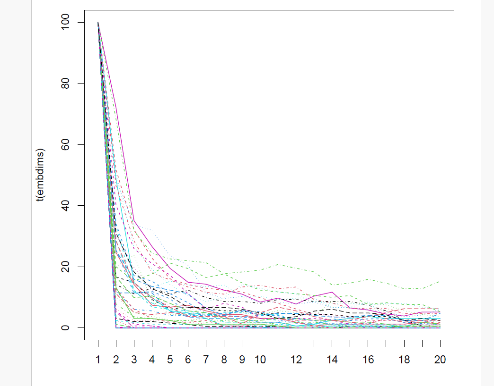 |
|  | **Selected *m* = 3** | | | |
| Radius | With *d* = 15 and *m* = 3, the selected radius was *r* = 1.114. The segment means were 7.816469, 3.501876, 3.937523, and 4.754945; their average was 5.002703% REC. | | | |

|  | IPS-RESP | | | |
| --- | --- | --- | --- | --- |
|  | BL | T1 | MAN | T2 |
| Delays | 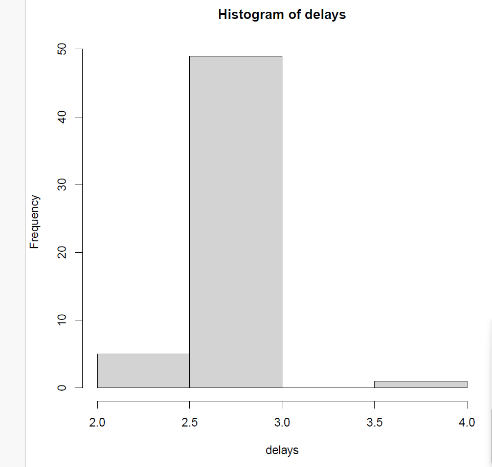 | 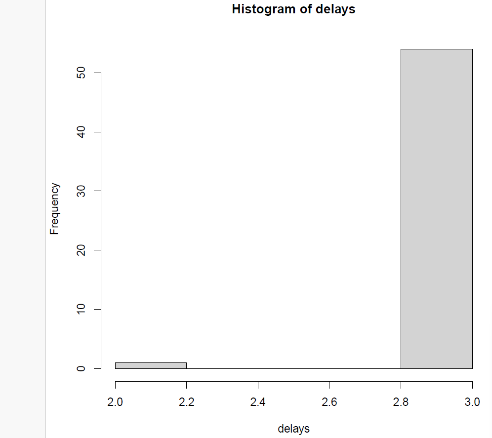 | 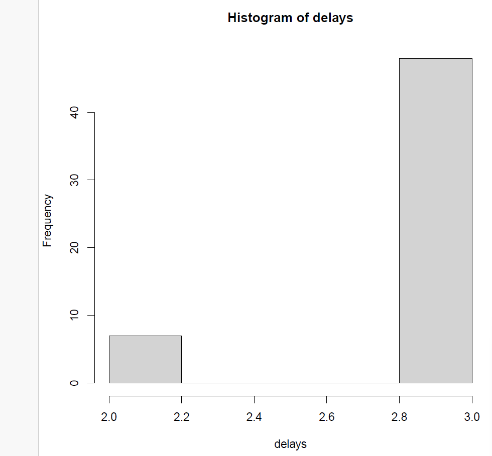 | 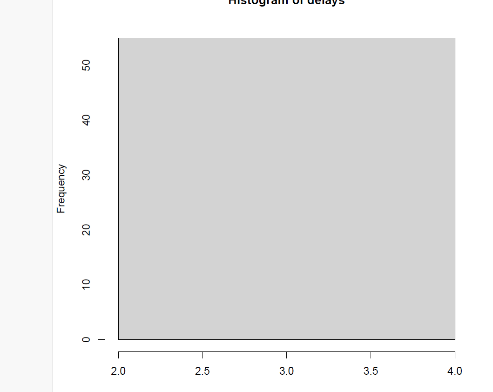 |
|  | Max = 4, mean = 2.927273, *SD* = 0.3251521 | Max = 3, mean = 2.981818, *SD* = 0.13484 | Max = 3, mean = 2.872727, *SD* = 0.33635 | Max = 3, mean = 3.000000, *SD* = 0 |
|  | Mean across segment means = 2.945454; **selected *d* = 3** | | | |
| False nearest neighbors | 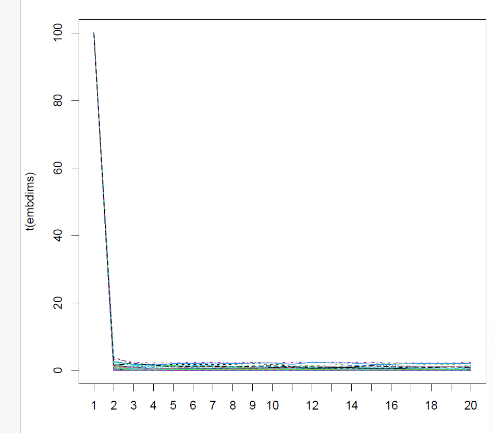 | 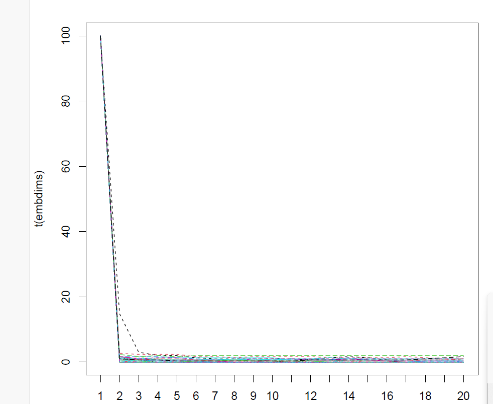 | 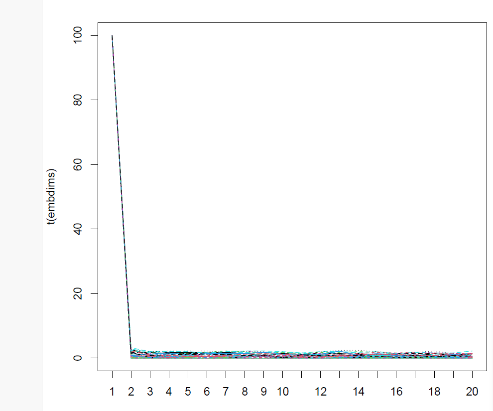 | 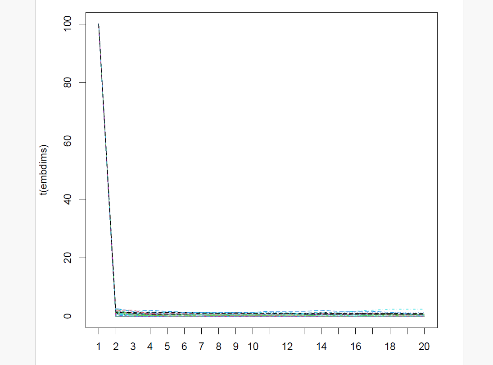 |
|  | **Selected *m* = 2** | | | |
| Radius | With *d* = 3 and *m* = 2, the selected radius was *r* = 1.325. The segment means were 5.478974, 3.666744, 6.707021, and 4.168989; their average was 5.005432% REC. | | | |

**Table S1**

*Gender Composition - Number of Women per Triad by ER Composition*

| ER composition | 0 women | 1 woman | 2 women | 3 women | Total |
| --- | --- | --- | --- | --- | --- |
| Condition 0 | 1 | 2 | 5 | 11 | 19 |
| Condition 1 | 1 | 4 | 6 | 5 | 16 |
| Condition 2 | 0 | 2 | 5 | 13 | 20 |
| Total | 2 | 8 | 16 | 29 | 55 |

*Note*. Triads were not deliberately matched or balanced by gender. A Pearson chi-square test was not run because expected-cell assumptions were inadequate: 7 of 12 expected counts (58.3%) were below 5 and the minimum expected count was .58. No alternative test was substituted.

**Table S2**

*Final-model R², Adjusted ICCs, and Random-effect and Residual Standard Deviations*

| Outcome | Marginal *R²* [95% CI] | Conditional *R²* [95% CI] | Adjusted ICC [95% CI] | Random-effect and residual SDs (and intercept-slope r) |
| --- | --- | --- | --- | --- |
| Participant IAPS | 0.038 [0.018, 0.072] | 0.183 [0.107, 0.265] | 0.151 [0.073, 0.232] | 0.71; 1.88; 4.77 |
| IPS-BPM | 0.260 [0.173, 0.387] | 0.535 [0.409, 0.663] | 0.371 [0.204, 0.522] | 1.39; 0.49; 2.50 |
| IPS-EDA | 0.250 [0.186, 0.347] | 0.418 [0.314, 0.537] | 0.224 [0.076, 0.363] | 1.58; 2.94 |
| IPS-RESP | 0.501 [0.403, 0.605] | 0.639 [0.559, 0.740] | 0.277 [0.133, 0.459] | 0.85; 0.28; *r* = .963; 1.83 |

*Note.* Confidence intervals for R² and adjusted ICCs were based on 1,000 parametric bootstrap iterations. The final column reports model-specific random-effect and residual quantities in the following order. For participant IAPS: triad-intercept SD, participant-intercept SD, and residual SD. For IPS-BPM: triad-intercept SD, within-triad IAPS-slope SD, and residual SD. For IPS-EDA: triad-intercept SD and residual SD. For IPS-RESP: triad-intercept SD, within-triad IAPS-slope SD, intercept-slope correlation (r), and residual SD. For models with random IAPS slopes, adjusted ICCs were calculated by averaging random-effect variance across the observed within-triad IAPS values. For each model, all 1,000 bootstrap draws yielded finite R² and ICC estimates. Boundary fits were retained and occurred in 281 participant-IAPS, 2 IPS-BPM, 1 IPS-EDA, and 543 IPS-RESP refits. The participant-IAPS and IPS-RESP bootstrap intervals, particularly those for IPS-RESP, should therefore be interpreted cautiously.

**Table S3**

*Mixed-effects Models Predicting IPS from Triad-average IAPS Across Segments, Separately Adjusted for Mean Heart Rate or Tonic Skin Conductance Level*

| Predictor | *b* | *SE* | *df* | *t* | *p* | 95% CI |
| --- | --- | --- | --- | --- | --- | --- |
| IPS-BPM | | | | | | |
| Intercept (grand mean; centered predictors) | 5.09 | 0.27 | 56.88 | 19.10 | < .001 | [4.56, 5.62] |
| IAPS (average slope; centered) | 0.58 | 0.09 | 105.70 | 6.66 | < .001 | [0.41, 0.76] |
| Segment: BL (sum contrast) | −1.13 | 0.34 | 175.17 | −3.34 | .001 | [−1.80, −0.46] |
| Segment: T1 (sum contrast) | 1.49 | 0.38 | 196.09 | 3.88 | < .001 | [0.73, 2.25] |
| Segment: MAN (sum contrast) | −0.87 | 0.34 | 183.57 | −2.55 | .012 | [−1.55, −0.20] |
| Mean heart rate (centered) | 0.03 | 0.04 | 67.74 | 0.84 | .406 | [−0.05, 0.12] |
| IAPS × BL (sum contrast) | 0.09 | 0.12 | 202.24 | 0.71 | .479 | [−0.15, 0.33] |
| IAPS × T1 (sum contrast) | 0.10 | 0.12 | 195.53 | 0.81 | .418 | [−0.14, 0.35] |
| IAPS × MAN (sum contrast) | −0.25 | 0.11 | 192.21 | −2.37 | .019 | [−0.46, −0.04] |
| IPS-EDA | | | | | | |
| Intercept (grand mean; centered predictors) | 5.13 | 0.29 | 58.33 | 17.52 | < .001 | [4.54, 5.71] |
| IAPS (average slope; centered) | 0.28 | 0.07 | 209.57 | 3.91 | < .001 | [0.14, 0.42] |
| Segment: BL (sum contrast) | 2.28 | 0.36 | 172.24 | 6.26 | < .001 | [1.56, 2.99] |
| Segment: T1 (sum contrast) | −0.85 | 0.39 | 170.69 | −2.22 | .028 | [−1.62, −0.09] |
| Segment: MAN (sum contrast) | −1.35 | 0.36 | 164.09 | −3.76 | < .001 | [−2.06, −0.64] |
| Tonic SCL (centered) | −0.20 | 0.07 | 59.55 | −2.82 | .006 | [−0.35, −0.06] |
| IAPS × BL (sum contrast) | −0.06 | 0.12 | 200.79 | −0.48 | .635 | [−0.29, 0.18] |
| IAPS × T1 (sum contrast) | 0.17 | 0.13 | 193.50 | 1.32 | .190 | [−0.09, 0.43] |
| IAPS × MAN (sum contrast) | −0.17 | 0.11 | 185.56 | −1.59 | .113 | [−0.38, 0.04] |

*Note*. Fixed effects are from models retaining the outcome-specific primary random structure: uncorrelated triad random intercepts and within-triad IAPS slopes for IPS-BPM and a triad random intercept for IPS-EDA. IAPS was centered and sum-to-zero contrasts were used for segment. The displayed BL, T1, and MAN terms are deviations from the average across segments; the T2 deviation is implied by their negative sum. The same interpretation applies to the displayed IAPS × segment contrasts.

**Table S4**

*Mixed-effects Models Predicting IPS from Triad-average IAPS Across Segments, Adjusted for RSA*

| Predictor | *b* | *SE* | *df* | *t* | *p* | 95% CI |
| --- | --- | --- | --- | --- | --- | --- |
| IPS-BPM | | | | | | |
| Intercept (grand mean; centered predictors) | 5.09 | 0.26 | 57.52 | 19.27 | < .001 | [4.56, 5.62] |
| IAPS (average slope; centered) | 0.57 | 0.09 | 106.61 | 6.47 | < .001 | [0.39, 0.74] |
| Segment: BL (sum contrast) | −1.23 | 0.32 | 155.08 | −3.80 | < .001 | [−1.87, −0.59] |
| Segment: T1 (sum contrast) | 1.66 | 0.36 | 170.89 | 4.60 | < .001 | [0.95, 2.37] |
| Segment: MAN (sum contrast) | −1.09 | 0.33 | 166.22 | −3.31 | .001 | [−1.75, −0.44] |
| RSA (centered) | −0.73 | 0.44 | 95.69 | −1.65 | .103 | [−1.60, 0.15] |
| IAPS × BL (sum contrast) | 0.09 | 0.12 | 203.39 | 0.75 | .452 | [−0.15, 0.33] |
| IAPS × T1 (sum contrast) | 0.11 | 0.12 | 196.06 | 0.87 | .386 | [−0.14, 0.35] |
| IAPS × MAN (sum contrast) | −0.25 | 0.11 | 192.81 | −2.40 | .017 | [−0.46, −0.05] |
| IPS-EDA | | | | | | |
| Intercept (grand mean; centered predictors) | 5.14 | 0.30 | 58.79 | 16.93 | < .001 | [4.53, 5.74] |
| IAPS (average slope; centered) | 0.30 | 0.07 | 208.50 | 4.04 | < .001 | [0.15, 0.44] |
| Segment: BL (sum contrast) | 2.47 | 0.36 | 161.52 | 6.81 | < .001 | [1.75, 3.18] |
| Segment: T1 (sum contrast) | −0.95 | 0.39 | 170.96 | −2.44 | .016 | [−1.73, −0.18] |
| Segment: MAN (sum contrast) | −1.30 | 0.37 | 176.69 | −3.47 | < .001 | [−2.04, −0.56] |
| RSA (centered) | 0.08 | 0.50 | 102.52 | 0.17 | .868 | [−0.91, 1.08] |
| IAPS × BL (sum contrast) | −0.06 | 0.12 | 199.43 | −0.52 | .605 | [−0.30, 0.17] |
| IAPS × T1 (sum contrast) | 0.18 | 0.13 | 192.30 | 1.34 | .182 | [−0.08, 0.44] |
| IAPS × MAN (sum contrast) | −0.18 | 0.11 | 184.90 | −1.70 | .090 | [−0.40, 0.03] |
| IPS-RESP | | | | | | |
| Intercept (grand mean; centered predictors) | 5.04 | 0.17 | 55.42 | 29.14 | < .001 | [4.69, 5.39] |
| IAPS (average slope; centered) | 0.53 | 0.06 | 91.03 | 8.66 | < .001 | [0.41, 0.65] |
| Segment: BL (sum contrast) | 0.10 | 0.23 | 156.76 | 0.41 | .684 | [−0.37, 0.56] |
| Segment: T1 (sum contrast) | −0.37 | 0.26 | 167.43 | −1.40 | .164 | [−0.88, 0.15] |
| Segment: MAN (sum contrast) | 0.90 | 0.24 | 168.12 | 3.81 | < .001 | [0.44, 1.37] |
| RSA (centered) | −0.00 | 0.28 | 87.12 | −0.00 | .998 | [−0.57, 0.56] |
| IAPS × BL (sum contrast) | −0.04 | 0.09 | 180.48 | −0.42 | .674 | [−0.21, 0.14] |
| IAPS × T1 (sum contrast) | 0.01 | 0.09 | 195.60 | 0.11 | .912 | [−0.17, 0.19] |
| IAPS × MAN (sum contrast) | 0.27 | 0.08 | 185.06 | 3.46 | < .001 | [0.12, 0.42] |

*Note.* Fixed effects are from models retaining the outcome-specific primary random structure: uncorrelated triad random intercepts and within-triad IAPS slopes for IPS-BPM, a triad random intercept for IPS-EDA, and correlated triad random intercepts and within-triad IAPS slopes for IPS-RESP. IAPS was centered and sum-to-zero contrasts were used for segment. The displayed BL, T1, and MAN terms are deviations from the average across segments; the T2 deviation is implied by their negative sum. The same interpretation applies to the displayed IAPS × segment contrasts.

**Table S5**

*Mixed-effects Models Predicting IPS from Triad-average IAPS Across Segments, Adjusted for Triad Mean Age*

| Predictor | *b* | *SE* | *df* | *t* | *p* | 95% CI |
| --- | --- | --- | --- | --- | --- | --- |
| IPS-BPM | | | | | | |
| Intercept (grand mean; centered predictors) | 5.10 | 0.26 | 58.81 | 19.48 | < .001 | [4.58, 5.63] |
| IAPS (average slope; centered) | 0.60 | 0.09 | 105.64 | 6.77 | < .001 | [0.42, 0.78] |
| Segment: BL (sum contrast) | −1.23 | 0.33 | 156.17 | −3.76 | < .001 | [−1.87, −0.58] |
| Segment: T1 (sum contrast) | 1.62 | 0.36 | 169.36 | 4.48 | < .001 | [0.91, 2.34] |
| Segment: MAN (sum contrast) | −0.98 | 0.32 | 154.74 | −3.04 | .003 | [−1.62, −0.34] |
| Triad mean age (centered) | −0.12 | 0.09 | 54.89 | −1.35 | .184 | [−0.29, 0.06] |
| IAPS × BL (sum contrast) | 0.07 | 0.12 | 202.56 | 0.60 | .547 | [−0.17, 0.32] |
| IAPS × T1 (sum contrast) | 0.11 | 0.12 | 197.14 | 0.89 | .375 | [−0.14, 0.36] |
| IAPS × MAN (sum contrast) | −0.26 | 0.11 | 193.96 | −2.45 | .015 | [−0.47, −0.05] |
| IPS-EDA | | | | | | |
| Intercept (grand mean; centered predictors) | 5.13 | 0.30 | 58.78 | 16.96 | < .001 | [4.53, 5.74] |
| IAPS (average slope; centered) | 0.29 | 0.07 | 207.52 | 3.90 | < .001 | [0.14, 0.43] |
| Segment: BL (sum contrast) | 2.47 | 0.36 | 161.74 | 6.83 | < .001 | [1.76, 3.19] |
| Segment: T1 (sum contrast) | −0.96 | 0.39 | 168.05 | −2.47 | .015 | [−1.73, −0.19] |
| Segment: MAN (sum contrast) | −1.30 | 0.36 | 163.84 | −3.59 | < .001 | [−2.02, −0.59] |
| Triad mean age (centered) | 0.08 | 0.10 | 54.98 | 0.77 | .446 | [−0.12, 0.28] |
| IAPS × BL (sum contrast) | −0.05 | 0.12 | 198.46 | −0.44 | .658 | [−0.29, 0.18] |
| IAPS × T1 (sum contrast) | 0.17 | 0.13 | 192.19 | 1.30 | .195 | [−0.09, 0.43] |
| IAPS × MAN (sum contrast) | −0.18 | 0.11 | 185.10 | −1.67 | .096 | [−0.39, 0.03] |
| IPS-RESP | | | | | | |
| Intercept (grand mean; centered predictors) | 5.03 | 0.17 | 56.88 | 29.11 | < .001 | [4.68, 5.37] |
| IAPS (average slope; centered) | 0.53 | 0.06 | 90.60 | 8.54 | < .001 | [0.41, 0.65] |
| Segment: BL (sum contrast) | 0.09 | 0.23 | 157.57 | 0.40 | .688 | [−0.37, 0.56] |
| Segment: T1 (sum contrast) | −0.37 | 0.26 | 165.59 | −1.43 | .155 | [−0.89, 0.14] |
| Segment: MAN (sum contrast) | 0.91 | 0.23 | 156.78 | 3.92 | < .001 | [0.45, 1.37] |
| Triad mean age (centered) | 0.03 | 0.05 | 56.63 | 0.52 | .607 | [−0.08, 0.13] |
| IAPS × BL (sum contrast) | −0.03 | 0.09 | 181.35 | −0.38 | .705 | [−0.21, 0.14] |
| IAPS × T1 (sum contrast) | 0.00 | 0.09 | 194.66 | 0.05 | .964 | [−0.17, 0.18] |
| IAPS × MAN (sum contrast) | 0.27 | 0.08 | 185.15 | 3.50 | < .001 | [0.12, 0.43] |

*Note*. Fixed effects are from models retaining the outcome-specific primary random structure: uncorrelated triad random intercepts and within-triad IAPS slopes for IPS-BPM, a triad random intercept for IPS-EDA, and correlated triad random intercepts and within-triad IAPS slopes for IPS-RESP. IAPS and triad mean age were centered, and sum-to-zero contrasts were used for segment. The displayed BL, T1, and MAN terms are deviations from the average across segments; the T2 deviation is implied by their negative sum. The same interpretation applies to the displayed IAPS × segment contrasts.

**Table S6**

*Mixed-effects Models Predicting IPS from Triad-average IAPS Across Segments, Adjusted for Pre-task Positive and Negative Affect*

| Predictor | *b* | *SE* | *df* | *t* | *p* | 95% CI |
| --- | --- | --- | --- | --- | --- | --- |
| IPS-BPM | | | | | | |
| Intercept (grand mean; centered predictors) | 5.09 | 0.27 | 56.99 | 19.17 | < .001 | [4.56, 5.63] |
| IAPS (average slope; centered) | 0.58 | 0.09 | 105.22 | 6.60 | < .001 | [0.41, 0.76] |
| Segment: BL (sum contrast) | −1.22 | 0.33 | 155.74 | −3.73 | < .001 | [−1.87, −0.57] |
| Segment: T1 (sum contrast) | 1.60 | 0.36 | 169.00 | 4.42 | < .001 | [0.89, 2.32] |
| Segment: MAN (sum contrast) | −0.96 | 0.32 | 154.44 | −2.98 | .003 | [−1.60, −0.32] |
| Pre-task positive affect (centered) | 0.53 | 0.71 | 51.50 | 0.75 | .456 | [−0.89, 1.96] |
| Pre-task negative affect (centered) | −0.59 | 0.91 | 51.28 | −0.64 | .523 | [−2.42, 1.25] |
| IAPS × BL (sum contrast) | 0.09 | 0.12 | 201.59 | 0.74 | .461 | [−0.15, 0.33] |
| IAPS × T1 (sum contrast) | 0.10 | 0.12 | 196.14 | 0.84 | .404 | [−0.14, 0.35] |
| IAPS × MAN (sum contrast) | −0.26 | 0.11 | 190.51 | −2.46 | .015 | [−0.48, −0.05] |
| IPS-EDA | | | | | | |
| Intercept (grand mean; centered predictors) | 5.14 | 0.31 | 57.36 | 16.77 | < .001 | [4.52, 5.75] |
| IAPS (average slope; centered) | 0.30 | 0.07 | 207.86 | 4.03 | < .001 | [0.15, 0.44] |
| Segment: BL (sum contrast) | 2.47 | 0.36 | 161.49 | 6.82 | < .001 | [1.75, 3.18] |
| Segment: T1 (sum contrast) | −0.95 | 0.39 | 168.12 | −2.43 | .016 | [−1.72, −0.18] |
| Segment: MAN (sum contrast) | −1.32 | 0.36 | 163.67 | −3.62 | < .001 | [−2.04, −0.60] |
| Pre-task positive affect (centered) | −0.32 | 0.82 | 52.59 | −0.39 | .695 | [−1.98, 1.33] |
| Pre-task negative affect (centered) | 0.06 | 1.06 | 52.27 | 0.06 | .951 | [−2.06, 2.19] |
| IAPS × BL (sum contrast) | −0.06 | 0.12 | 197.78 | −0.53 | .595 | [−0.30, 0.17] |
| IAPS × T1 (sum contrast) | 0.18 | 0.13 | 191.69 | 1.33 | .185 | [−0.08, 0.44] |
| IAPS × MAN (sum contrast) | −0.18 | 0.11 | 182.21 | −1.65 | .100 | [−0.39, 0.03] |
| IPS-RESP | | | | | | |
| Intercept (grand mean; centered predictors) | 4.88 | 0.17 | 54.77 | 28.45 | < .001 | [4.53, 5.22] |
| IAPS (average slope; centered) | 0.57 | 0.06 | 100.25 | 9.81 | < .001 | [0.46, 0.69] |
| Segment: BL (sum contrast) | 0.07 | 0.24 | 154.92 | 0.31 | .761 | [−0.40, 0.55] |
| Segment: T1 (sum contrast) | −0.39 | 0.26 | 168.90 | −1.50 | .134 | [−0.91, 0.12] |
| Segment: MAN (sum contrast) | 0.93 | 0.24 | 154.33 | 3.91 | < .001 | [0.46, 1.40] |
| Pre-task positive affect (centered) | −0.23 | 0.46 | 48.85 | −0.50 | .622 | [−1.14, 0.69] |
| Pre-task negative affect (centered) | 0.74 | 0.59 | 48.50 | 1.26 | .214 | [−0.44, 1.91] |
| IAPS × BL (sum contrast) | 0.00 | 0.09 | 194.14 | 0.03 | .980 | [−0.17, 0.17] |
| IAPS × T1 (sum contrast) | −0.05 | 0.09 | 201.72 | −0.53 | .599 | [−0.22, 0.13] |
| IAPS × MAN (sum contrast) | 0.31 | 0.08 | 193.56 | 4.05 | < .001 | [0.16, 0.46] |

*Note*. Pre-task positive and negative affect were averaged within triad and centered. IPS-BPM retained uncorrelated triad random intercepts and within-triad IAPS slopes, and IPS-EDA retained a triad random intercept. The correlated IPS-RESP structure was singular after adding both affect covariates, so the documented hierarchy selected the corresponding uncorrelated intercept-and-slope structure. IAPS was centered and sum-to-zero contrasts were used for segment.

**Table S7**

*Mixed-effects Models Predicting IPS from Triad-average IAPS Across Segments, with ER Composition as an Additive Control*

| Predictor | *b* | *SE* | *df* | *t* | *p* | 95% CI |
| --- | --- | --- | --- | --- | --- | --- |
| IPS-BPM | | | | | | |
| Intercept (grand mean; centered predictors) | 5.05 | 0.26 | 57.83 | 19.06 | < .001 | [4.52, 5.58] |
| IAPS (average slope; centered) | 0.60 | 0.09 | 103.03 | 6.77 | < .001 | [0.43, 0.78] |
| Segment: BL (sum contrast) | −1.22 | 0.33 | 155.75 | −3.74 | < .001 | [−1.87, −0.58] |
| Segment: T1 (sum contrast) | 1.61 | 0.36 | 169.22 | 4.44 | < .001 | [0.89, 2.32] |
| Segment: MAN (sum contrast) | −0.97 | 0.32 | 154.48 | −3.00 | .003 | [−1.61, −0.33] |
| ER composition: contrast 1 | 0.01 | 0.36 | 52.29 | 0.04 | .971 | [−0.71, 0.73] |
| ER composition: contrast 2 | −0.47 | 0.38 | 54.60 | −1.24 | .220 | [−1.24, 0.29] |
| IAPS × BL (sum contrast) | 0.10 | 0.12 | 200.67 | 0.82 | .413 | [−0.14, 0.34] |
| IAPS × T1 (sum contrast) | 0.09 | 0.13 | 195.49 | 0.73 | .464 | [−0.16, 0.34] |
| IAPS × MAN (sum contrast) | −0.25 | 0.11 | 193.18 | −2.30 | .022 | [−0.46, −0.03] |
| IPS-EDA | | | | | | |
| Intercept (grand mean; centered predictors) | 5.17 | 0.30 | 57.99 | 17.19 | < .001 | [4.56, 5.77] |
| IAPS (average slope; centered) | 0.29 | 0.07 | 207.30 | 3.91 | < .001 | [0.14, 0.43] |
| Segment: BL (sum contrast) | 2.48 | 0.36 | 161.46 | 6.86 | < .001 | [1.77, 3.20] |
| Segment: T1 (sum contrast) | −0.95 | 0.39 | 168.38 | −2.44 | .016 | [−1.72, −0.18] |
| Segment: MAN (sum contrast) | −1.32 | 0.36 | 163.88 | −3.63 | < .001 | [−2.04, −0.60] |
| ER composition: contrast 1 | −0.71 | 0.41 | 52.93 | −1.74 | .088 | [−1.53, 0.11] |
| ER composition: contrast 2 | 0.43 | 0.43 | 54.49 | 0.99 | .325 | [−0.44, 1.29] |
| IAPS × BL (sum contrast) | −0.08 | 0.12 | 198.12 | −0.70 | .485 | [−0.32, 0.15] |
| IAPS × T1 (sum contrast) | 0.19 | 0.13 | 191.93 | 1.45 | .150 | [−0.07, 0.45] |
| IAPS × MAN (sum contrast) | −0.18 | 0.11 | 185.13 | −1.68 | .095 | [−0.39, 0.03] |
| IPS-RESP | | | | | | |
| Intercept (grand mean; centered predictors) | 5.05 | 0.17 | 58.21 | 29.27 | < .001 | [4.71, 5.40] |
| IAPS (average slope; centered) | 0.52 | 0.06 | 90.37 | 8.33 | < .001 | [0.39, 0.64] |
| Segment: BL (sum contrast) | 0.10 | 0.23 | 158.90 | 0.45 | .655 | [−0.36, 0.57] |
| Segment: T1 (sum contrast) | −0.37 | 0.26 | 167.03 | −1.43 | .154 | [−0.89, 0.14] |
| Segment: MAN (sum contrast) | 0.92 | 0.23 | 158.37 | 3.96 | < .001 | [0.46, 1.37] |
| ER composition: contrast 1 | −0.04 | 0.22 | 56.59 | −0.18 | .854 | [−0.49, 0.41] |
| ER composition: contrast 2 | 0.29 | 0.23 | 57.52 | 1.26 | .212 | [−0.17, 0.76] |
| IAPS × BL (sum contrast) | −0.04 | 0.09 | 179.63 | −0.50 | .618 | [−0.22, 0.13] |
| IAPS × T1 (sum contrast) | 0.02 | 0.09 | 193.45 | 0.23 | .818 | [−0.16, 0.20] |
| IAPS × MAN (sum contrast) | 0.26 | 0.08 | 185.40 | 3.39 | < .001 | [0.11, 0.42] |

*Note*. ER composition was the three-level variable indicating number of members with elevated ER-difficulties. The displayed coefficients are sum-to-zero contrasts. ER composition was included additively in this secondary robustness model; no IAPS × ER composition moderation term was fitted. The models retained the outcome-specific primary random structures: an uncorrelated triad intercept-and-IAPS-slope structure for IPS-BPM, a triad random intercept for IPS-EDA, and a correlated triad intercept-and-IAPS-slope structure for IPS-RESP. IAPS was centered and sum-to-zero contrasts were used for segment.

**Table S8**

*Candidate Random-effects Structures and Selection*

| Analysis | Candidate structure | Fitted | Converged | Singular | Selected | Reason |
| --- | --- | --- | --- | --- | --- | --- |
| Participant IAPS | correlated segment slopes at both levels | No | No | N/A | No | Unidentifiable: observations did not exceed random effects |
| Participant IAPS | uncorrelated segment slopes at both levels | Yes | Yes | Yes | No | Singular fit |
| Participant IAPS | participant segment slopes only | Yes | Yes | Yes | No | Singular fit |
| Participant IAPS | triad segment slopes only | Yes | Yes | Yes | No | Singular fit |
| Participant IAPS | triad and participant random intercepts | Yes | Yes | No | Yes | First supported candidate in the hierarchy |
| IPS-BPM | correlated segment slopes | No | No | N/A | No | Unidentifiable: observations did not exceed random effects |
| IPS-BPM | uncorrelated segment slopes | Yes | Yes | Yes | No | Singular fit |
| IPS-BPM | correlated within-triad IAPS slope | Yes | Yes | Yes | No | Singular fit |
| IPS-BPM | uncorrelated within-triad IAPS slope | Yes | Yes | No | Yes | First supported candidate in the hierarchy |
| IPS-BPM | triad random intercept | Yes | Yes | No | No | Supported but lower in the hierarchy |
| IPS-EDA | correlated segment slopes | No | No | N/A | No | Unidentifiable: observations did not exceed random effects |
| IPS-EDA | uncorrelated segment slopes | Yes | Yes | Yes | No | Singular fit |
| IPS-EDA | correlated within-triad IAPS slope | Yes | Yes | Yes | No | Singular fit |
| IPS-EDA | uncorrelated within-triad IAPS slope | Yes | Yes | Yes | No | Singular fit |
| IPS-EDA | triad random intercept | Yes | Yes | No | Yes | First supported candidate in the hierarchy |
| IPS-RESP | correlated segment slopes | No | No | N/A | No | Unidentifiable: observations did not exceed random effects |
| IPS-RESP | uncorrelated segment slopes | Yes | Yes | Yes | No | Singular fit |
| IPS-RESP | correlated within-triad IAPS slope | Yes | Yes | No | Yes | Converged, nonsingular, and optimizer-stable |
| IPS-RESP | uncorrelated within-triad IAPS slope | Yes | Yes | No | No | Supported but lower in the hierarchy |
| IPS-RESP | triad random intercept | Yes | Yes | No | No | Supported but lower in the hierarchy |

*Note*. The corresponding maximal-to-simple hierarchy was evaluated for each outcome, and the first identifiable, converged, nonsingular candidate was retained. A correlated within-triad IAPS-slope candidate additionally had to be stable across optimizers. The selected participant IAPS model retained triad and participant random intercepts; IPS-BPM retained uncorrelated triad random intercepts and within-triad IAPS slopes; IPS-EDA retained a triad random intercept; and IPS-RESP retained correlated triad random intercepts and within-triad IAPS slopes. The RESP intercept-slope correlation was high (*r* = .963), but the model remained nonsingular and stable across three optimizers. An uncorrelated RESP sensitivity retained the positive average IAPS effect and a significant interaction. The Holm-adjusted MAN-versus-T2 contrast was significant in both models, whereas the Holm-adjusted T1-versus-MAN contrast was significant only in the uncorrelated sensitivity. AIC and BIC were descriptive and were not used to select among nonnested structures.

**Table S9a**

*Formal Residual Checks and Segment-specific Residual-variance Comparisons*

| Outcome | Raw residual-normality *p* | Raw heteroscedasticity *p* | LR | VarIdent *p* | Log1p residual-normality *p* | Log1p heteroscedasticity *p* |
| --- | --- | --- | --- | --- | --- | --- |
| Participant IAPS | < .001 | < .001 | 23.93 | < .001 | < .001 | .274 |
| IPS-BPM | < .001 | < .001 | 19.24 | < .001 | .002 | .694 |
| IPS-EDA | < .001 | < .001 | 81.21 | < .001 | .042 | .821 |
| IPS-RESP | < .001 | < .001 | 18.79 | < .001 | < .001 | .425 |

*Note*. The likelihood-ratio comparison evaluated a common residual variance against residual variances allowed to differ across segments. Formal residual-normality checks remained significant after log1p transformation, whereas heteroscedasticity was no longer detected. Diagnostic plots are shown in Figures S1–S3.

**Table S9b**

*Omnibus Tests Across Primary and Sensitivity Specifications*

| Outcome | Effect | Primary raw scale | Segment-specific residual variance | Log1p outcome | Within-between |
| --- | --- | --- | --- | --- | --- |
| Participant IAPS | Segment | *F* = 10.28, *p* < .001 | *F* = 11.66, *p* < .001 | *F* = 18.53, *p* < .001 | Not applicable |
| IPS-BPM | Average IAPS slope | *F* = 43.91, *p* < .001 | *F* = 49.84, *p* < .001 | *F* = 66.56, *p* < .001 | *F* = 35.32, *p* < .001 |
| IPS-BPM | IAPS × segment | *F* = 1.97, *p* = .120 | *F* = 2.08, *p* = .104 | *F* = 0.47, *p* = .706 | *F* = 1.24, *p* = .298 |
| IPS-EDA | Average IAPS slope | *F* = 16.39, *p* < .001 | *F* = 16.25, *p* < .001 | *F* = 36.56, *p* < .001 | *F* = 8.84, *p* = .003 |
| IPS-EDA | IAPS × segment | *F* = 1.28, *p* = .284 | *F* = 3.08, *p* = .029 | *F* = 3.30, *p* = .021 | *F* = 1.07, *p* = .364 |
| IPS-RESP | Average IAPS slope | *F* = 76.06, *p* < .001 | *F* = 68.58, *p* < .001 | *F* = 111.11, *p* < .001 | *F* = 35.80, *p* < .001 |
| IPS-RESP | IAPS × segment | *F* = 4.73, *p* = .003 | *F* = 6.42, *p* < .001 | *F* = 1.52, *p* = .212 | *F* = 3.94, *p* = .009 |

*Note*. The within-between average IAPS entry is the within-triad IAPS effect; each separate between-triad effect is reported in Table S11. The EDA interaction was supported in the unequal-variance and log1p models but not in the primary or within-between models. The RESP interaction was supported in the primary, unequal-variance, and within-between models but not in the log1p model. All four passive–active segment contrasts remained significant after Holm correction in both the segment-specific residual-variance and log1p sensitivity models. The T1–T2 contrast was additionally significant only in the log1p model (Holm-adjusted *p* = .006; primary Holm-adjusted *p* = .133; segment-specific residual-variance Holm-adjusted *p* = .060).

**Table S9c**

*Segment-specific IAPS Slopes Across Model Specifications*

| Outcome | Specification | BL | T1 | MAN | T2 |
| --- | --- | --- | --- | --- | --- |
| IPS-BPM | Primary raw scale | 0.671 [0.379, 0.963] | 0.682 [0.373, 0.991] | 0.325 [0.073, 0.577] | 0.637 [0.309, 0.966] |
| IPS-BPM | Segment-specific residual variance | 0.667 [0.301, 1.032] | 0.692 [0.360, 1.025] | 0.355 [0.172, 0.537] | 0.621 [0.347, 0.895] |
| IPS-BPM | Log1p outcome | 0.078 [0.044, 0.111] | 0.086 [0.049, 0.122] | 0.064 [0.035, 0.093] | 0.087 [0.050, 0.125] |
| IPS-BPM | Within-triad IAPS | 0.669 [0.320, 1.019] | 0.871 [0.483, 1.258] | 0.402 [0.033, 0.772] | 0.730 [0.337, 1.123] |
| IPS-EDA | Primary raw scale | 0.233 [−0.033, 0.500] | 0.473 [0.155, 0.790] | 0.111 [−0.127, 0.350] | 0.364 [0.079, 0.648] |
| IPS-EDA | Segment-specific residual variance | 0.217 [−0.219, 0.652] | 0.454 [0.291, 0.617] | 0.087 [−0.102, 0.276] | 0.367 [0.166, 0.569] |
| IPS-EDA | Log1p outcome | 0.055 [0.020, 0.090] | 0.101 [0.059, 0.143] | 0.020 [−0.012, 0.051] | 0.059 [0.022, 0.097] |
| IPS-EDA | Within-triad IAPS | 0.337 [0.018, 0.656] | 0.353 [−0.022, 0.728] | −0.037 [−0.386, 0.312] | 0.314 [−0.031, 0.658] |
| IPS-RESP | Primary raw scale | 0.496 [0.285, 0.706] | 0.543 [0.325, 0.762] | 0.804 [0.622, 0.986] | 0.291 [0.054, 0.529] |
| IPS-RESP | Segment-specific residual variance | 0.488 [0.260, 0.716] | 0.599 [0.445, 0.753] | 0.799 [0.589, 1.009] | 0.167 [−0.070, 0.404] |
| IPS-RESP | Log1p outcome | 0.069 [0.045, 0.094] | 0.075 [0.048, 0.103] | 0.090 [0.068, 0.111] | 0.054 [0.026, 0.081] |
| IPS-RESP | Within-triad IAPS | 0.442 [0.201, 0.683] | 0.311 [0.042, 0.581] | 0.816 [0.558, 1.074] | 0.277 [0.006, 0.547] |

*Note*. Entries are slope [95% CI]. Log1p slopes are on the transformed outcome scale. Within-between slopes represent the association between a triad's segment-specific deviation from its own mean IAPS and IPS. No BPM slope contrast survived Holm correction. For EDA, only T1 versus MAN survived in the unequal-variance (*p* = .020) and log1p (*p* = .013) models. For RESP, only MAN versus T2 survived in the primary model (*p* = .003); T1 versus T2 and MAN versus T2 survived in the unequal-variance model (*p* = .004 and *p* < .001); MAN versus T1 and MAN versus T2 survived in the within-between model (*p* = .022 and *p* = .017); and no comparison survived in the log1p model.

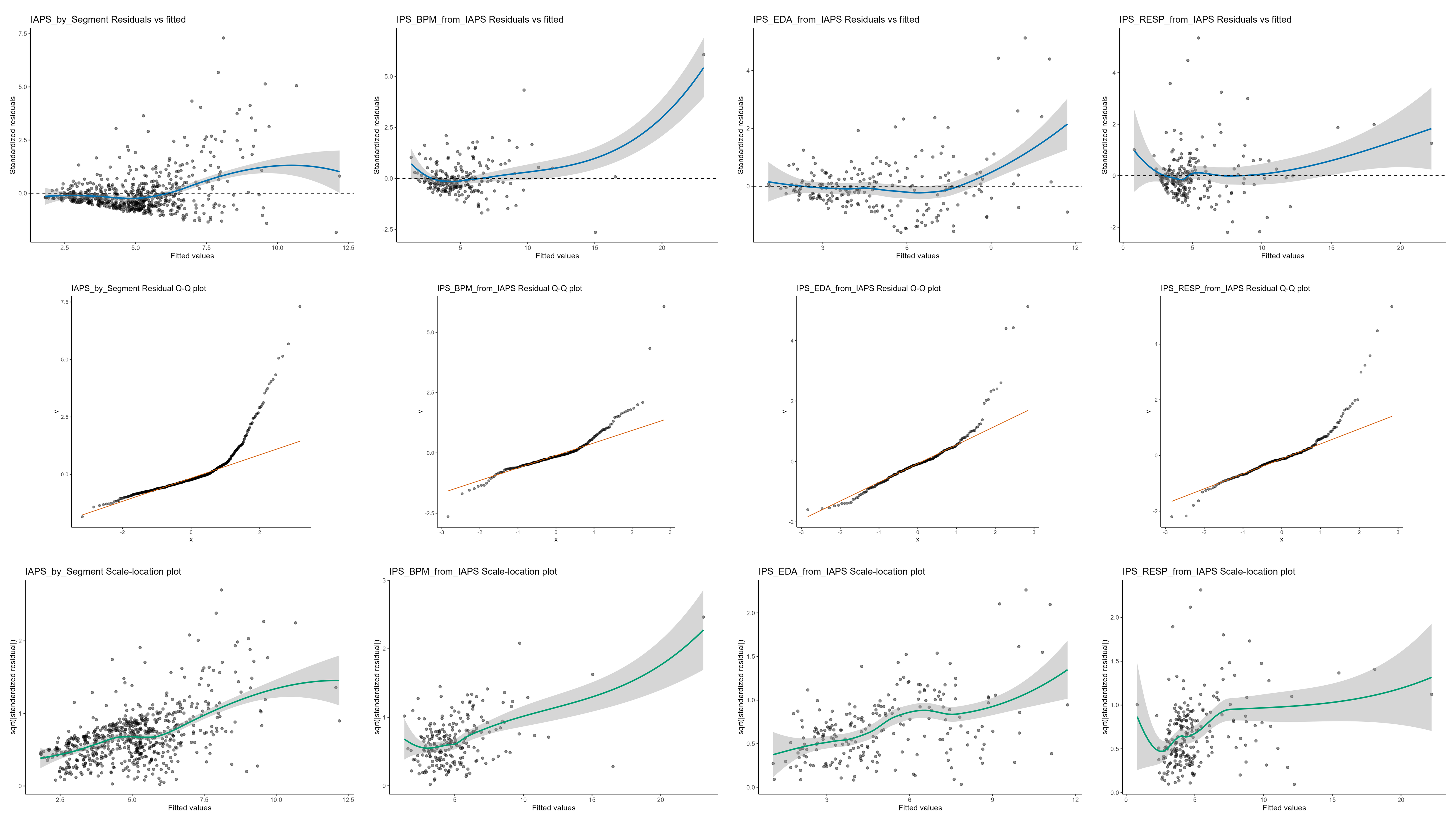

**Figure S1.** Diagnostic plots for the primary raw-scale mixed models. Columns show participant-level IAPS, IPS-BPM, IPS-EDA, and IPS-RESP; rows show residuals versus fitted values, residual Q-Q plots, and scale-location plots.

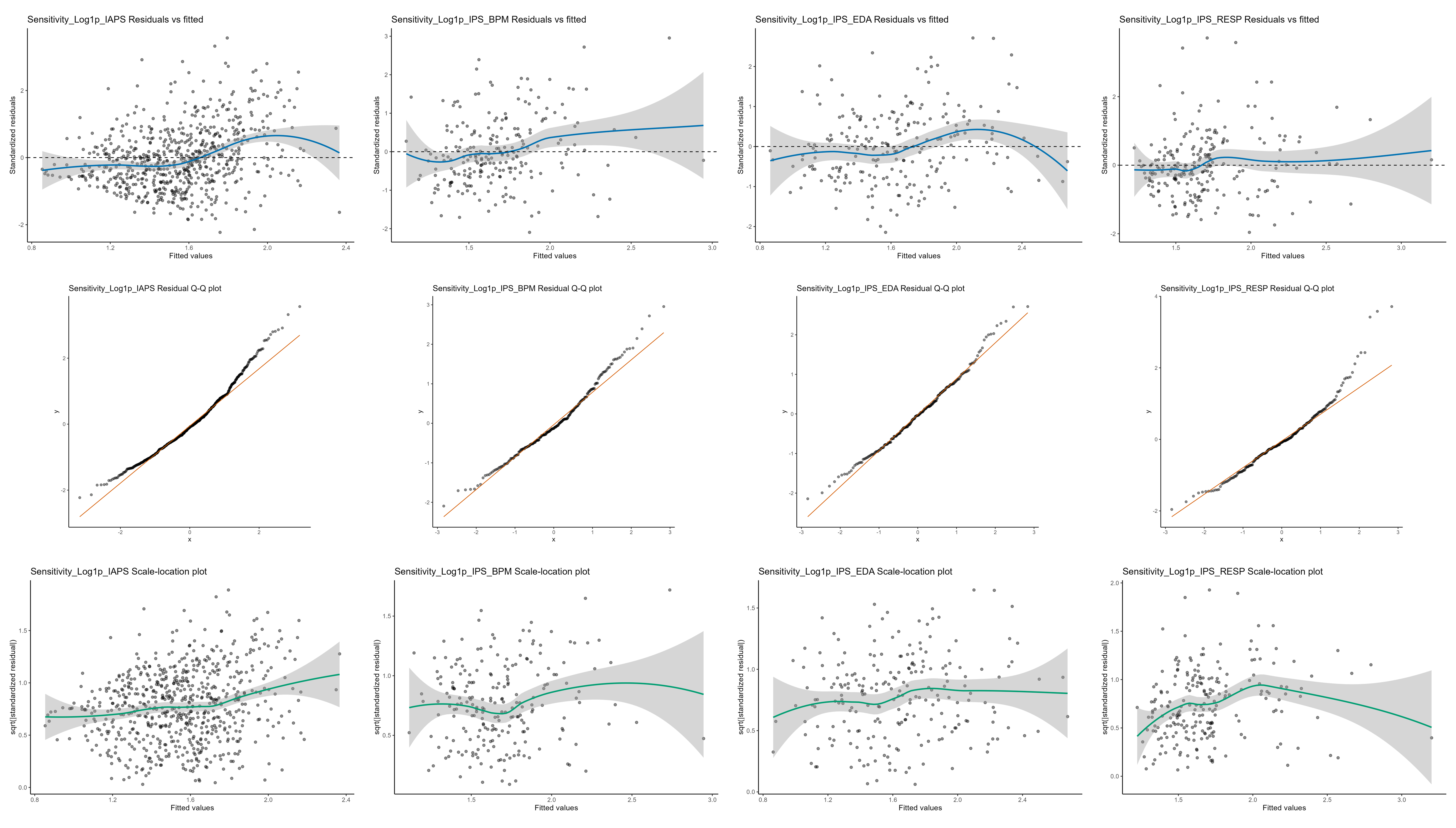

**Figure S2.** Diagnostic plots for the corresponding log1p sensitivity models. Columns show participant-level IAPS, IPS-BPM, IPS-EDA, and IPS-RESP; rows show residuals versus fitted values, residual Q-Q plots, and scale-location plots.

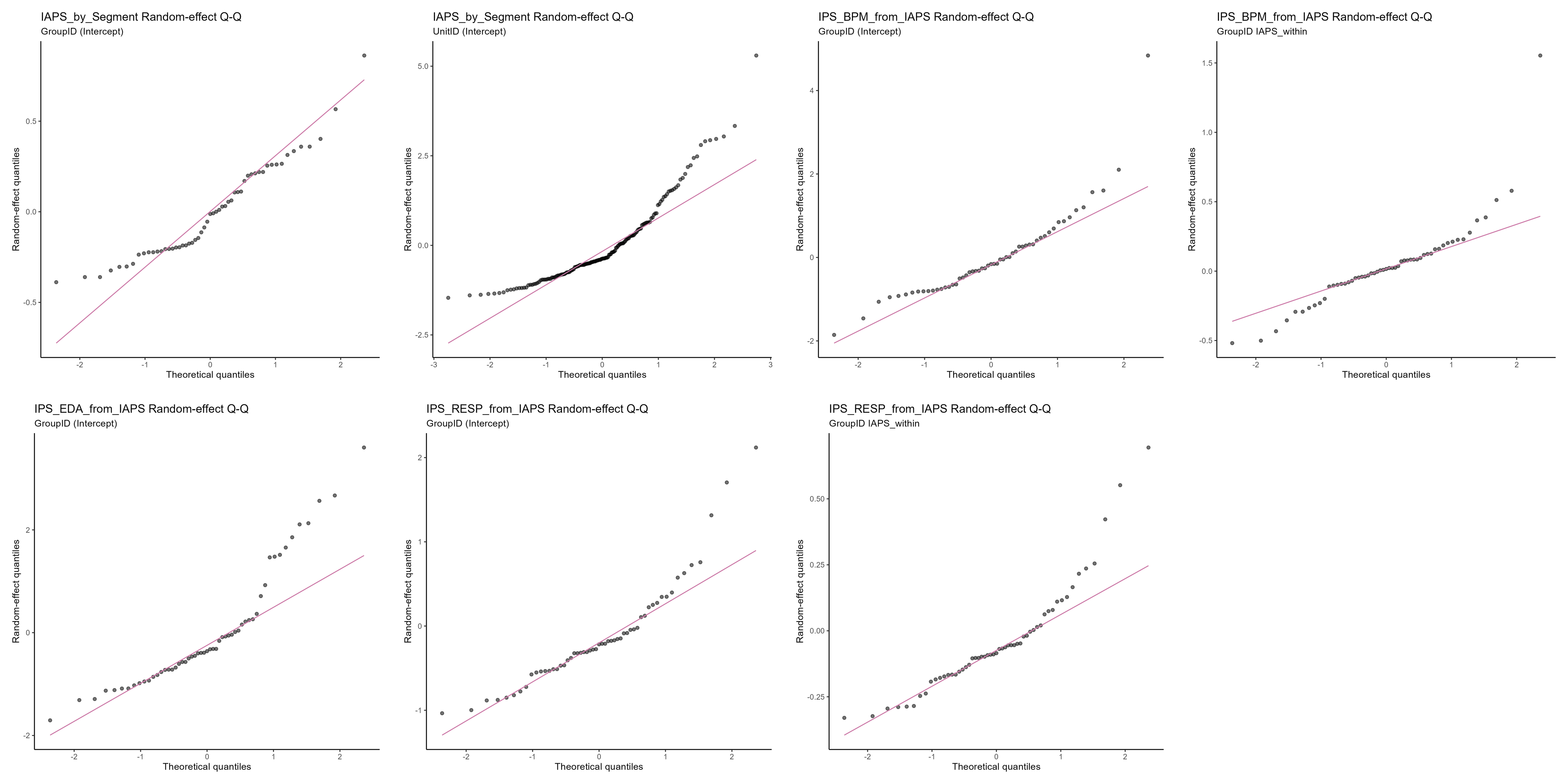

**Figure S3.** Random-effect Q-Q plots for the primary models. Panels show the triad and participant random intercepts for participant-level IAPS; the triad random intercept and within-triad IAPS slope for IPS-BPM; the triad random intercept for IPS-EDA; and the triad random intercept and within-triad IAPS slope for IPS-RESP.

**Table S10**

*Triad-level Influence and Leave-one-triad-out Results*

| Analysis | Valid/attempted refits | Maximum Cook's D | Cook's D flags | DFBETAS flags | Focal omnibus p range | Interaction p range |
| --- | --- | --- | --- | --- | --- | --- |
| Participant IAPS | 55/55 | 0.115 | 3 | 11 | < .001 to < .001 | Not applicable |
| IPS-BPM | 55/55 | 1.044 | 4 | 12 | < .001 to < .001 | .043 to .173 |
| IPS-EDA | 55/55 | 0.147 | 4 | 12 | < .001 to < .001 | .077 to .396 |
| IPS-RESP | 53/55 | 0.599 | 7 | 14 | < .001 to < .001 | < .001 to .028 |

*Note*. Thresholds were 4/55 for Cook's D and 2/√55 for absolute DFBETAS. All 55 IAPS, BPM, and EDA deletion refits were valid. For RESP, all 55 deletion refits returned, and 53 were valid for inference. One returned fit was nonconverged and two were singular; because these statuses overlapped, two unique refits were invalid. Reported ranges therefore summarize valid refits only and may understate sensitivity across all deletions. The positive average IAPS association remained significant in every valid BPM, EDA, and RESP refit, and the RESP IAPS × segment interaction remained significant in all 53 valid refits. For participant-level IAPS, the BL–T1, MAN–T1, and MAN–T2 contrasts remained Holm-significant in all 55 deletion refits; BL–T2 remained Holm-significant in 51 of 55. Influence flags prompted sensitivity assessment; no triad was excluded solely because it was flagged.

**Table S11**

*Within-between Decomposition Models Predicting IPS*

| Predictor | *b* | *SE* | *df* | *t* | *p* | 95% CI |
| --- | --- | --- | --- | --- | --- | --- |
| IPS-BPM | | | | | | |
| Intercept (grand mean; centered predictors) | 5.15 | 0.27 | 60.40 | 19.14 | < .001 | [4.61, 5.69] |
| IAPS between triads (centered) | 0.44 | 0.14 | 53.41 | 3.25 | .002 | [0.17, 0.72] |
| IAPS within triad | 0.67 | 0.11 | 55.74 | 5.94 | < .001 | [0.44, 0.89] |
| Segment: BL (sum contrast) | −1.26 | 0.33 | 156.26 | −3.76 | < .001 | [−1.92, −0.60] |
| Segment: T1 (sum contrast) | 1.75 | 0.38 | 170.94 | 4.63 | < .001 | [1.00, 2.49] |
| Segment: MAN (sum contrast) | −1.01 | 0.33 | 150.45 | −3.02 | .003 | [−1.67, −0.35] |
| Within-triad IAPS × BL | 0.00 | 0.15 | 208.81 | 0.01 | .993 | [−0.29, 0.29] |
| Within-triad IAPS × T1 | 0.20 | 0.15 | 207.88 | 1.33 | .186 | [−0.10, 0.50] |
| Within-triad IAPS × MAN | −0.27 | 0.15 | 210.45 | −1.75 | .082 | [−0.57, 0.03] |
| IPS-EDA | | | | | | |
| Intercept (grand mean; centered predictors) | 5.10 | 0.31 | 61.10 | 16.45 | < .001 | [4.48, 5.72] |
| IAPS between triads (centered) | 0.40 | 0.16 | 54.24 | 2.55 | .014 | [0.09, 0.72] |
| IAPS within triad | 0.24 | 0.08 | 159.66 | 2.97 | .003 | [0.08, 0.40] |
| Segment: BL (sum contrast) | 2.41 | 0.37 | 162.32 | 6.50 | < .001 | [1.68, 3.14] |
| Segment: T1 (sum contrast) | −1.09 | 0.40 | 167.01 | −2.69 | .008 | [−1.88, −0.29] |
| Segment: MAN (sum contrast) | −1.13 | 0.38 | 164.13 | −2.98 | .003 | [−1.87, −0.38] |
| Within-triad IAPS × BL | 0.10 | 0.15 | 209.69 | 0.64 | .521 | [−0.20, 0.39] |
| Within-triad IAPS × T1 | 0.11 | 0.16 | 205.89 | 0.68 | .496 | [−0.21, 0.43] |
| Within-triad IAPS × MAN | −0.28 | 0.16 | 206.66 | −1.77 | .079 | [−0.59, 0.03] |
| IPS-RESP | | | | | | |
| Intercept (grand mean; centered predictors) | 4.84 | 0.17 | 62.21 | 28.80 | < .001 | [4.51, 5.18] |
| IAPS between triads (centered) | 0.74 | 0.08 | 54.65 | 8.80 | < .001 | [0.57, 0.91] |
| IAPS within triad | 0.46 | 0.08 | 54.26 | 5.98 | < .001 | [0.31, 0.62] |
| Segment: BL (sum contrast) | 0.16 | 0.25 | 159.50 | 0.65 | .517 | [−0.33, 0.65] |
| Segment: T1 (sum contrast) | −0.59 | 0.27 | 174.24 | −2.14 | .033 | [−1.13, −0.05] |
| Segment: MAN (sum contrast) | 0.93 | 0.25 | 154.89 | 3.76 | < .001 | [0.44, 1.42] |
| Within-triad IAPS × BL | −0.02 | 0.10 | 196.11 | −0.19 | .849 | [−0.22, 0.18] |
| Within-triad IAPS × T1 | −0.15 | 0.11 | 210.90 | −1.39 | .165 | [−0.36, 0.06] |
| Within-triad IAPS × MAN | 0.35 | 0.11 | 208.39 | 3.32 | .001 | [0.14, 0.56] |

*Note*. IAPS between triads is each triad's mean IAPS across segments, centered at the sample mean. IAPS within triad is each segment's deviation from that triad's own mean. Thus, the between effect compares triads with different average IAPS, whereas the within effect tests whether IPS changes when the same triad's IAPS is above or below its own average. Segment used sum-to-zero contrasts. BPM retained the uncorrelated intercept-and-IAPS-slope structure and EDA retained a random intercept. The correlated RESP structure was singular in the expanded within-between model, so the documented hierarchy selected the corresponding uncorrelated intercept-and-slope structure.

**Table S12**

*Parametric Simulation-Refit Design Sensitivity for Average IAPS Slopes*

| Target | Standardized β | Valid/requested | All-requested detection | Exact 95% CI |
| --- | --- | --- | --- | --- |
| IPS-BPM average IAPS slope | 0 | 1000/1000 | 0.080 | [0.064, 0.099] |
| IPS-BPM average IAPS slope | 0.05 | 1000/1000 | 0.098 | [0.080, 0.118] |
| IPS-BPM average IAPS slope | 0.1 | 1000/1000 | 0.254 | [0.227, 0.282] |
| IPS-BPM average IAPS slope | 0.15 | 1000/1000 | 0.500 | [0.469, 0.531] |
| IPS-BPM average IAPS slope | 0.2 | 1000/1000 | 0.705 | [0.676, 0.733] |
| IPS-BPM average IAPS slope | 0.25 | 1000/1000 | 0.876 | [0.854, 0.896] |
| IPS-BPM average IAPS slope | 0.3 | 1000/1000 | 0.970 | [0.957, 0.980] |
| IPS-EDA average IAPS slope | 0 | 1000/1000 | 0.050 | [0.037, 0.065] |
| IPS-EDA average IAPS slope | 0.05 | 1000/1000 | 0.139 | [0.118, 0.162] |
| IPS-EDA average IAPS slope | 0.1 | 1000/1000 | 0.349 | [0.319, 0.379] |
| IPS-EDA average IAPS slope | 0.15 | 1000/1000 | 0.676 | [0.646, 0.705] |
| IPS-EDA average IAPS slope | 0.2 | 1000/1000 | 0.899 | [0.879, 0.917] |
| IPS-EDA average IAPS slope | 0.25 | 1000/1000 | 0.975 | [0.963, 0.984] |
| IPS-RESP average IAPS slope | 0 | 960/1000 | 0.058 | [0.044, 0.074] |
| IPS-RESP average IAPS slope | 0.05 | 969/1000 | 0.127 | [0.107, 0.149] |
| IPS-RESP average IAPS slope | 0.1 | 977/1000 | 0.345 | [0.316, 0.375] |
| IPS-RESP average IAPS slope | 0.15 | 968/1000 | 0.693 | [0.663, 0.721] |
| IPS-RESP average IAPS slope | 0.2 | 966/1000 | 0.860 | [0.837, 0.881] |
| IPS-RESP average IAPS slope | 0.25 | 971/1000 | 0.948 | [0.932, 0.961] |

*Note*. Each grid point requested 1,000 simulations. Detection required a two-sided Satterthwaite-adjusted *p* < .05. The all-requested rate uses 1,000 as its denominator, so failed refits do not inflate the estimate. Operational designation required at least 80% all-requested detection at two consecutive positive grid points, at least 95% valid refits, and an exact beta-zero binomial interval containing α = .05. IPS-EDA met these criteria at β = .20 (detection = .899). IPS-RESP met them at β = .20 (detection = .860; 966 valid refits), but 520 of its 966 finite refits were singular, so this estimate is approximate. IPS-BPM reached .876 detection at β = .25 but was not designated because its beta-zero rejection rate was .080, 95% CI [.064, .099].

**Table S13**

*IAPS Comparisons Below Versus At or Above Clinical or Screening Cutoffs*

| Measure | Cutoff | Segment | M below | M at/above | *t* | *df* | *p* | Hedges' *g* | pHolm |
| --- | --- | --- | --- | --- | --- | --- | --- | --- | --- |
| DERS | 95 | BL | 5.72 | 6.27 | 0.55 | 90.40 | .585 | 0.093 | 1.000 |
| DERS | 95 | T1 | 3.23 | 4.13 | 1.14 | 75.00 | .257 | 0.200 | 1.000 |
| DERS | 95 | MAN | 6.39 | 5.36 | −1.18 | 147.09 | .240 | −0.183 | 1.000 |
| DERS | 95 | T2 | 4.50 | 4.53 | 0.03 | 115.23 | .975 | 0.005 | 1.000 |
| SPIN | 19 | BL | 6.00 | 5.85 | −0.17 | 149.88 | .863 | −0.027 | 1.000 |
| SPIN | 19 | T1 | 3.38 | 3.66 | 0.44 | 144.74 | .662 | 0.069 | 1.000 |
| SPIN | 19 | MAN | 6.67 | 5.60 | −1.17 | 144.50 | .245 | −0.184 | 1.000 |
| SPIN | 19 | T2 | 4.63 | 4.43 | −0.23 | 113.16 | .817 | −0.038 | 1.000 |
| GAD-7 | 10 | BL | 5.90 | 6.07 | 0.13 | 42.34 | .894 | 0.028 | 1.000 |
| GAD-7 | 10 | T1 | 3.40 | 4.16 | 0.75 | 40.60 | .460 | 0.159 | 1.000 |
| GAD-7 | 10 | MAN | 6.18 | 5.60 | −0.56 | 56.59 | .581 | −0.103 | 1.000 |
| GAD-7 | 10 | T2 | 4.51 | 4.44 | −0.06 | 45.06 | .950 | −0.013 | 1.000 |
| PHQ-2 | 3 | BL | 5.65 | 6.72 | 0.93 | 60.26 | .354 | 0.175 | 1.000 |
| PHQ-2 | 3 | T1 | 3.56 | 3.56 | 0.01 | 79.70 | .996 | 0.001 | 1.000 |
| PHQ-2 | 3 | MAN | 6.52 | 4.81 | −1.93 | 106.11 | .056 | −0.313 | .841 |
| PHQ-2 | 3 | T2 | 4.86 | 3.49 | −2.19 | 157.94 | .030 | −0.317 | .478 |

*Note*. Welch independent-samples *t* tests compared participants at or above each cutoff with participants below it. Test statistics and Hedges' *g* are coded as at/above minus below. Holm correction was applied across all 16 comparisons. No comparison remained significant after correction. In an auxiliary triad-clustered sensitivity, all 12 estimable tests remained nonsignificant after Holm correction; the four baseline models were singular and were not interpreted.

**Table S14**

*Recorded ER-Composition Condition as a Predictor of Participant-Level IAPS*

| Effect or contrast | Estimate | *SE* | *df* | *F* or *t* | *p* | Partial *R²* | 95% CI |
| --- | --- | --- | --- | --- | --- | --- | --- |
| **Panel A** | | | | | | | |
| Segment |  |  | 3, 486.00 | 11.14 | < .001 | 0.064 | [0.025, 0.107] |
| ER composition |  |  | 2, 52.00 | 3.32 | .044 | 0.113 | [0.000, 0.275] |
| Segment × ER composition |  |  | 6, 486.00 | 1.70 | .119 | 0.021 | [0.000, 0.040] |
| **Panel B** | | | | | | | |
| Condition 0 | 4.68 | 0.42 | 52.00 |  |  |  | [3.84, 5.51] |
| Condition 1 | 5.98 | 0.45 | 52.00 |  |  |  | [5.07, 6.89] |
| Condition 2 | 4.52 | 0.41 | 52.00 |  |  |  | [3.71, 5.34] |
| **Panel C** | | | | | | | |
| Condition 0 − Condition 1 | −1.30 | 0.62 | 52.00 | −2.11 | .079 |  | [−2.82, 0.22] |
| Condition 0 − Condition 2 | 0.16 | 0.58 | 52.00 | 0.27 | .790 |  | [−1.28, 1.59] |
| Condition 1 − Condition 2 | 1.46 | 0.61 | 52.00 | 2.39 | .061 |  | [−0.05, 2.96] |

*Note*. Panel A gives omnibus Type III tests. Panel B gives estimated marginal means averaged across the four segments. Panel C gives Holm-adjusted *p* values and Bonferroni-adjusted simultaneous 95% confidence intervals for pairwise comparisons. The recorded parent-study Condition variable was used for these analyses. Conditions 0, 1, and 2 denote triads with zero, one, and two members assigned to the low-ER recruitment category, respectively, and contained 19, 16, and 20 triads. Partial *R²* values were derived from the omnibus *F* tests.

The following provides full statistics for the high–low synchrony analyses reported in the main manuscript. Analyses involving overall performance used 54 complete cases because one triad lacked complete overall-performance data; physiological profile definitions and group-efficacy analyses retained all 55 analytic triads. Holm correction was applied across the four BPM and EDA omnibus Welch tests. As an exploratory supplementary extension, the same profile procedure was repeated using IPS-RESP; the two RESP tests are reported with unadjusted *p* values and were not added to that four-test family.

### **Table S15a**

### *Descriptive Statistics for Group Outcomes by High–Low Synchrony Classification*

| Composite | Outcome | Subgroup | *N* | Mean | *SD* | *SE* |
| --- | --- | --- | --- | --- | --- | --- |
| IAPS × IPS-EDA | Overall performance | High–High | 21 | 29.14 | 7.39 | 1.613 |
|  |  | High–Low | 7 | 19.43 | 11.70 | 4.423 |
|  |  | Low–High | 7 | 19.57 | 5.80 | 2.192 |
|  |  | Low–Low | 19 | 24.79 | 10.86 | 2.490 |
| IAPS × IPS-EDA | Group efficacy | High–High | 21 | 4.34 | 0.76 | 0.166 |
|  |  | High–Low | 7 | 4.27 | 0.87 | 0.331 |
|  |  | Low–High | 7 | 4.34 | 0.35 | 0.132 |
|  |  | Low–Low | 20 | 4.35 | 0.77 | 0.172 |
| IAPS × IPS-BPM | Overall performance | High–High | 20 | 27.30 | 9.04 | 2.022 |
|  |  | High–Low | 8 | 25.25 | 10.90 | 3.853 |
|  |  | Low–High | 8 | 25.62 | 8.93 | 3.156 |
|  |  | Low–Low | 18 | 22.39 | 10.42 | 2.456 |
| IAPS × IPS-BPM | Group efficacy | High–High | 20 | 4.54 | 0.77 | 0.171 |
|  |  | High–Low | 8 | 3.77 | 0.48 | 0.170 |
|  |  | Low–High | 8 | 4.50 | 0.31 | 0.109 |
|  |  | Low–Low | 19 | 4.29 | 0.78 | 0.180 |
| IAPS × IPS-RESP | Overall performance | High–High | 22 | 28.55 | 9.59 | 2.045 |
|  |  | High–Low | 6 | 20.00 | 5.18 | 2.113 |
|  |  | Low–High | 6 | 19.67 | 8.91 | 3.639 |
|  |  | Low–Low | 20 | 24.50 | 10.15 | 2.269 |
| IAPS × IPS-RESP | Group efficacy | High–High | 22 | 4.29 | 0.84 | 0.180 |
|  |  | High–Low | 6 | 4.45 | 0.48 | 0.196 |
|  |  | Low–High | 6 | 4.73 | 0.57 | 0.234 |
|  |  | Low–Low | 21 | 4.24 | 0.68 | 0.148 |

### **Table S15b**

### *Welch One-Way ANOVA Results for High–Low Synchrony Composites*

| Composite | Outcome | *F* | df1 | df2 | *p (Holm-adjusted p)* | *η² (descriptive)* |
| --- | --- | --- | --- | --- | --- | --- |
| IAPS × IPS-EDA | Overall performance | 4.43 | 3 | 17.42 | .017; .052 | 0.156 |
| IAPS × IPS-EDA | Group efficacy | 0.02 | 3 | 19.38 | .997; 1.000 | 0.001 |
| IAPS × IPS-BPM | Overall performance | 0.75 | 3 | 19.49 | .538; 1.000 | 0.046 |
| IAPS × IPS-BPM | Group efficacy | 4.75 | 3 | 23.88 | .010; .039 | 0.131 |
| IAPS × IPS-RESP | Overall performance | 3.08 | 3 | 16.27 | .057; not adjusted | 0.120 |
| IAPS × IPS-RESP | Group efficacy | 1.06 | 3 | 15.98 | .394; not adjusted | 0.044 |

### **Table S15c**

### *Games–Howell Post Hoc Comparisons (All Pairwise Contrasts) with Hedges’ g*

| Composite | Outcome | Group 1 | Group 2 | Mean diff. (1–2) | *p (adj.)* | *Hedges’ g* |
| --- | --- | --- | --- | --- | --- | --- |
| IAPS × IPS-EDA | Overall performance | High–High | High–Low | 9.71 | .246 | 0.964 |
| IAPS × IPS-EDA | Overall performance | High–High | Low–High | 9.57 | .017 | 1.399 |
| IAPS × IPS-EDA | Overall performance | High–High | Low–Low | 4.35 | .469 | 0.459 |
| IAPS × IPS-EDA | Overall performance | High–Low | Low–High | −0.14 | 1.000 | −0.014 |
| IAPS × IPS-EDA | Overall performance | High–Low | Low–Low | −5.36 | .722 | −0.460 |
| IAPS × IPS-EDA | Overall performance | Low–High | Low–Low | −5.22 | .415 | −0.581 |
| IAPS × IPS-EDA | Group efficacy | High–High | High–Low | 0.07 | .998 | 0.082 |
| IAPS × IPS-EDA | Group efficacy | High–High | Low–High | 0.00 | 1.000 | 0.005 |
| IAPS × IPS-EDA | Group efficacy | High–High | Low–Low | −0.02 | 1.000 | −0.021 |
| IAPS × IPS-EDA | Group efficacy | High–Low | Low–High | −0.07 | .998 | −0.092 |
| IAPS × IPS-EDA | Group efficacy | High–Low | Low–Low | −0.08 | .996 | −0.100 |
| IAPS × IPS-EDA | Group efficacy | Low–High | Low–Low | −0.02 | 1.000 | −0.031 |
| IAPS × IPS-BPM | Overall performance | High–High | High–Low | 2.05 | .964 | 0.199 |
| IAPS × IPS-BPM | Overall performance | High–High | Low–High | 1.68 | .969 | 0.181 |
| IAPS × IPS-BPM | Overall performance | High–High | Low–Low | 4.91 | .424 | 0.493 |
| IAPS × IPS-BPM | Overall performance | High–Low | Low–High | −0.38 | 1.000 | −0.036 |
| IAPS × IPS-BPM | Overall performance | High–Low | Low–Low | 2.86 | .922 | 0.260 |
| IAPS × IPS-BPM | Overall performance | Low–High | Low–Low | 3.24 | .849 | 0.323 |
| IAPS × IPS-BPM | Group efficacy | High–High | High–Low | 0.77 | .021* | 1.176 |
| IAPS × IPS-BPM | Group efficacy | High–High | Low–High | 0.04 | .997 | 0.068 |
| IAPS × IPS-BPM | Group efficacy | High–High | Low–Low | 0.26 | .731 | 0.325 |
| IAPS × IPS-BPM | Group efficacy | High–Low | Low–High | −0.73 | .016* | −1.720 |
| IAPS × IPS-BPM | Group efficacy | High–Low | Low–Low | −0.52 | .189 | −0.771 |
| IAPS × IPS-BPM | Group efficacy | Low–High | Low–Low | 0.22 | .734 | 0.352 |
| IAPS × IPS-RESP | Overall performance | High–High | High–Low | 8.55 | .047 | 1.077 |
| IAPS × IPS-RESP | Overall performance | High–High | Low–High | 8.88 | .220 | 0.931 |
| IAPS × IPS-RESP | Overall performance | High–High | Low–Low | 4.05 | .553 | 0.402 |
| IAPS × IPS-RESP | Overall performance | High–Low | Low–High | 0.33 | 1.000 | 0.042 |
| IAPS × IPS-RESP | Overall performance | High–Low | Low–Low | −4.50 | .486 | −0.541 |
| IAPS × IPS-RESP | Overall performance | Low–High | Low–Low | −4.83 | .683 | −0.490 |
| IAPS × IPS-RESP | Group efficacy | High–High | High–Low | −0.17 | .924 | −0.234 |
| IAPS × IPS-RESP | Group efficacy | High–High | Low–High | −0.44 | .473 | −0.593 |
| IAPS × IPS-RESP | Group efficacy | High–High | Low–Low | 0.04 | .998 | 0.056 |
| IAPS × IPS-RESP | Group efficacy | High–Low | Low–High | −0.27 | .806 | −0.479 |
| IAPS × IPS-RESP | Group efficacy | High–Low | Low–Low | 0.21 | .830 | 0.344 |
| IAPS × IPS-RESP | Group efficacy | Low–High | Low–Low | 0.48 | .355 | 0.745 |

*Note.* Welch one-way ANOVAs accommodated unequal group sizes and heterogeneous variances. Holm correction was applied across the four BPM and EDA omnibus tests. The two IPS-RESP omnibus tests were exploratory and are reported without multiplicity adjustment. Games–Howell *p* values adjust the six pairwise comparisons within each analysis. Pairwise Hedges’ *g* values used an unpooled standardizer with small-sample correction. Mean differences and g are coded as Group 1 minus Group 2. Asterisks require both the Holm-adjusted omnibus *p* value and the Games–Howell-adjusted pairwise *p* value to be below .05. The exploratory RESP omnibus tests were nonsignificant, so no RESP pairwise comparison is interpreted. Diff. = difference; adj. = adjusted.

**Table S16**

*Approximate Pairwise Minimum Detectable Hedges’ g Values for High–Low Profile Comparisons*

| Analysis | Pair | n1, n2 | Variance ratio | MDE \|g\| at α = .05 | MDE \|g\| at α = .05/6 |
| --- | --- | --- | --- | --- | --- |
| Performance, BPM profiles | High–High vs High–Low | 20, 8 | 1.45 | 1.30 | 1.75 |
| Performance, BPM profiles | High–High vs Low–High | 20, 8 | 1.03 | 1.23 | 1.63 |
| Performance, BPM profiles | High–High vs Low–Low | 20, 18 | 1.33 | 0.92 | 1.17 |
| Performance, BPM profiles | High–Low vs Low–High | 8, 8 | 1.49 | 1.43 | 1.89 |
| Performance, BPM profiles | High–Low vs Low–Low | 8, 18 | 1.09 | 1.26 | 1.67 |
| Performance, BPM profiles | Low–High vs Low–Low | 8, 18 | 1.36 | 1.19 | 1.57 |
| Performance, EDA profiles | High–High vs High–Low | 21, 7 | 2.51 | 1.50 | 2.12 |
| Performance, EDA profiles | High–High vs Low–High | 21, 7 | 1.62 | 1.20 | 1.60 |
| Performance, EDA profiles | High–High vs Low–Low | 21, 19 | 2.16 | 0.91 | 1.15 |
| Performance, EDA profiles | High–Low vs Low–High | 7, 7 | 4.07 | 1.58 | 2.18 |
| Performance, EDA profiles | High–Low vs Low–Low | 7, 19 | 1.16 | 1.35 | 1.84 |
| Performance, EDA profiles | Low–High vs Low–Low | 7, 19 | 3.51 | 1.09 | 1.41 |
| Efficacy, BPM profiles | High–High vs High–Low | 20, 8 | 2.54 | 1.08 | 1.39 |
| Efficacy, BPM profiles | High–High vs Low–High | 20, 8 | 6.21 | 0.98 | 1.26 |
| Efficacy, BPM profiles | High–High vs Low–Low | 20, 19 | 1.05 | 0.90 | 1.15 |
| Efficacy, BPM profiles | High–Low vs Low–High | 8, 8 | 2.44 | 1.44 | 1.93 |
| Efficacy, BPM profiles | High–Low vs Low–Low | 8, 19 | 2.67 | 1.08 | 1.40 |
| Efficacy, BPM profiles | Low–High vs Low–Low | 8, 19 | 6.52 | 1.00 | 1.28 |
| Efficacy, EDA profiles | High–High vs High–Low | 21, 7 | 1.32 | 1.38 | 1.89 |
| Efficacy, EDA profiles | High–High vs Low–High | 21, 7 | 4.71 | 1.02 | 1.31 |
| Efficacy, EDA profiles | High–High vs Low–Low | 21, 20 | 1.02 | 0.88 | 1.12 |
| Efficacy, EDA profiles | High–Low vs Low–High | 7, 7 | 6.24 | 1.61 | 2.25 |
| Efficacy, EDA profiles | High–Low vs Low–Low | 7, 20 | 1.29 | 1.37 | 1.88 |
| Efficacy, EDA profiles | Low–High vs Low–Low | 7, 20 | 4.83 | 1.03 | 1.33 |
| Performance, RESP profiles | High–High vs High–Low | 22, 6 | 3.43 | 1.11 | 1.46 |
| Performance, RESP profiles | High–High vs Low–High | 22, 6 | 1.16 | 1.39 | 1.93 |
| Performance, RESP profiles | High–High vs Low–Low | 22, 20 | 1.12 | 0.87 | 1.11 |
| Performance, RESP profiles | High–Low vs Low–High | 6, 6 | 2.97 | 1.71 | 2.38 |
| Performance, RESP profiles | High–Low vs Low–Low | 6, 20 | 3.84 | 1.11 | 1.45 |
| Performance, RESP profiles | Low–High vs Low–Low | 6, 20 | 1.30 | 1.36 | 1.87 |
| Efficacy, RESP profiles | High–High vs High–Low | 22, 6 | 3.07 | 1.13 | 1.49 |
| Efficacy, RESP profiles | High–High vs Low–High | 22, 6 | 2.16 | 1.22 | 1.63 |
| Efficacy, RESP profiles | High–High vs Low–Low | 22, 21 | 1.53 | 0.86 | 1.09 |
| Efficacy, RESP profiles | High–Low vs Low–High | 6, 6 | 1.42 | 1.66 | 2.27 |
| Efficacy, RESP profiles | High–Low vs Low–Low | 6, 21 | 2.01 | 1.24 | 1.67 |
| Efficacy, RESP profiles | Low–High vs Low–Low | 6, 21 | 1.41 | 1.34 | 1.83 |

*Note*. Values are Welch-based pairwise approximations using the observed cell sizes and variance ratios, with target power of 80%. Performance rows use outcome-specific complete cases, whereas efficacy rows use all available analytic triads. The α = .05/6 column is a conservative six-comparison benchmark. For BPM and EDA, MDE |g| values ranged from 0.88 to 1.61 at α = .05 and from 1.12 to 2.25 at α = .05/6. For the exploratory RESP extension, they ranged from 0.86 to 1.71 and from 1.09 to 2.38, respectively. These values describe design sensitivity and do not establish equivalence when an observed comparison is nonsignificant.
